# Detection of histone modifications and full-length transcriptome in single cells identifies sequential epigenetic perceptiveness to lineage commitment

**DOI:** 10.1101/2024.05.09.593364

**Authors:** Marloes Blotenburg, Vivek Bhardwaj, Buys Anton de Barbanson, Fredrik Salmén, Peter Zeller, Alexander van Oudenaarden

## Abstract

Epigenetic mechanisms, including histone modifications, are key regulators of transcription and influence cellular differentiation. To uncover the interplay between histone modifications and transcription, we developed T-ChIC (Transcriptome + Chromatin ImmunoCleavage), which detects full-length transcripts and histone modifications in the same single cell. We applied T-ChIC to gastruloids, an in vitro model containing cell types derived from all three germ layers.

During gastruloid development we observed a switch in transcription start site usage from multiple regions marked by H3K4me3 to one marked start site per gene. H3K27me3 coverage transitions from global marking of non-transcribed regions towards localised marking of repressive regions.

Upon lineage commitment, bivalent regions are sequentially resolved towards a germ layer-specific state, where endoderm retains a high abundance of H3K27me3 while ectoderm displays a widespread loss. Together, we propose a model for a cell-intrinsic epigenetic timer, which ensures sequential perceptiveness of pluripotent cells to transcriptional activation of germ layer-specific genes.

## Introduction

During multicellular organism development, individual cells that share the same origin and genome need to specify towards different lineages at the correct time and location to form the complex mature body plan. Underlying this immense diversification is the ability of different cell types to transcribe distinct parts of their genome, resulting in cell type-specific gene expression patterns. In order for genes to become activated, their promoter region needs to be open and accessible for gene transcription factors to bind. A central regulator of genome accessibility are epigenetic modifications to the tail of histone proteins that bind and compact the DNA in a chromatin complex. These histone post-translational modifications (HPTM) are positioned and removed by mark-specific writer and eraser complexes respectively, that together shape cell type-specific epigenomes. Mutants of multiple writer complex subunits were shown to cause lineage-specific defects ^1–4^ or embryonic lethality at different timepoints ^5^. The Polycomb and Trithorax group proteins deposit H3K27me3 and H3K4me3 respectively, and their HPTM products play fundamental roles through their opposing function. While H3K4me3 marks active gene promoters and plays an essential role in PolII-mediated transcription ^6^, H3K27me3 represses gene expression. This opposing function is exemplified at the Hox gene cluster, which encodes a cell’s position along the anteroposterior axis of the developing embryo ^7–10^. This locus is regulated through the progressive removal of the H3K27me3 and deposition of H3K4me3 along the gene cluster ^8,10^. When the distribution of these two chromatin modifications along the Hox gene cluster is altered, cells adapt either a more posterior or anterior identity.

While the epigenome during pre-implantation development is extensively studied, epigenetic changes in post-implantation embryos are less characterised. On the onset of gastrulation, the embryo undergoes vast rearrangements and greatly increases in size and complexity, with precursors of all cell types rapidly emerging ^11^. Whereas pre-implantation stages only consist of a few different cell types and can be profiled using bulk technologies, the gastrula-stage embryo is highly complex. Recently, the gastruloid system has been developed to facilitate the study of gastrula-stage lineage specifications *in vitro*. This model mimics key aspects of gastrulation, displays temporal-spatial gene regulation, collinear Hox gene expression, and contains cell types derived from all three germ layers ^12–17^. Recent advances in single-cell RNA sequencing have allowed a detailed understanding of cell type complexity and dynamics in these developing systems, ^18–20^. However, our knowledge regarding epigenetic changes underlying gene regulation during lineage specification still remain largely unexplored.

Recently, a variety of techniques have been developed to study chromatin modifications at the single-cell level. Most of these are based on antibody tethering of proteinA-Tn5 transposase fusion (pA-Tn5) ^21–23^ or proteinA-micrococcal nuclease (pA-MN), such as sort-assisted chromatin immunocleavage (sortChIC), ^24–27^. However, to study the interplay between transcription and HPTMs and to tease apart cell type-specific profiles, co-acquisition of the cell’s nascent transcriptome is necessary. To this end, several pA-Tn5 based multi-omics approaches have been developed which recover both a HPTM and transcriptome from the same single cell ^28–30^. While some of these approaches allow the processing of large cell numbers by applying combinatorial barcoding ^29,30^ or 10x microfluidics ^28^, they only capture the 3’-end of transcripts. This largely inhibits the ability to profile unspliced nascent RNA, which most closely reflects current transcriptional activity. As a result, the temporal disconnect between accumulated spliced RNA and active transcription hinders accurate understanding of the interplay between chromatin regulation and active transcription.

Here, we developed T-ChIC (Transcriptome + Chromatin ImmunoCleavage), a novel multi-omic method to profile full-length transcriptome and HPTMs in the same single cell. This method combines VASA-seq ^31^ for profiling of the full-length transcriptome with sortChIC as a read-out of chromatin modifications. We demonstrate that T-ChIC provides a highly sensitive and specific read-out of genome-wide chromatin modification distributions in combination with the full transcriptome. We apply this technique to an *in vitro* model of gastrulation to study the epigenetic changes underlying lineage specification and provide cell type-specific characterisations of chromatin states during differentiation. Last, we provide a model for differential H3K27me3-mediated de-repression upon differentiation and sequential perceptiveness to lineage specification.

## Results

### T-ChIC profiles the full transcriptome and HPTM distribution in the same single cell with high sensitivity and specificity

In order to measure the transcriptome and HPTM distribution in the same single cell, we developed T-ChIC (Transcriptome + Chromatin Immunocleavage) which integrates sortChIC for profiling HPTMs ^27^ with VASA-seq for characterisation of the full-length transcriptome ^31^.To detect both signals in single cells, cells are fixed in ethanol, incubated with an antibody against the HPTM of interest and pA-MNase fusion is added that binds to the HPTM-bound antibody. G1-phase single cells are sorted into individual wells of 384-well plates, followed by pA-MNase activation, inactivation and cell lysis, resulting in double-strand breaks around the HPTM of interest, which are end repaired and ligated to adapters with a unique molecular identifier (UMI) and ChIC and cell-specific barcode. For full-transcript coverage, the RNA is fragmented, repaired and poly-A-tailed, followed by reverse transcription with primers containing a UMI, a transcript- and a cell-specific barcode. Plates are pooled, second strands are synthesised and products are amplified through in vitro transcription and PCR before sequencing of final libraries (Figure 1A). The ChIC and transcript-specific cell barcodes are used to computationally distinguish both signals in each cell.

**Figure 1.**
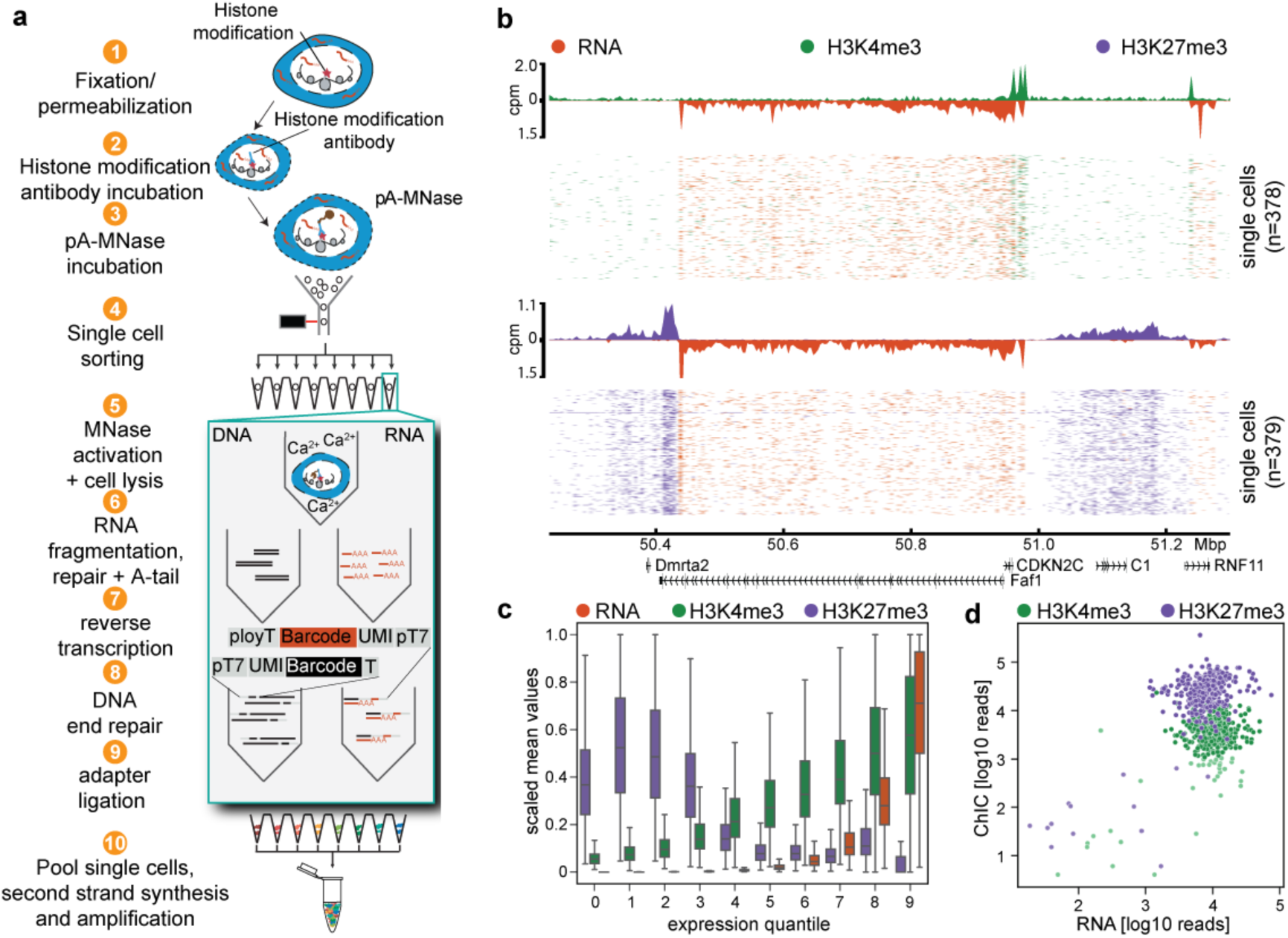
T-ChIC allows the simultaneous recovery of full-length transcript and HPTM position from the same single cell. a) T-ChIC experimental workflow including cell fixation and pA-MN targeting in bulk (1-3), sorting of single cells into 384-well plates by FACS (4), followed by separate processing of RNA and DNA fragments inside the plate and introducing RNA- and DNA-specific cell barcodes (5-9). After pooling, fragments are linearly amplified by IVT before sequencing handles are introduced in a final indexing PCR (10). b) Single-cell coverage plots from K562 cells processed for transcriptome (orange) and H3K4me3 (top, green) or H3K27me3 (bottom, purple) over a 1 Mb region of chromosome 2. Coverage plots above represent the average signals across cells. c) Boxplots of single-cell read averages over genes split into quantiles of log10 expression. RNA (orange), H3K4me3 (green) and H3K27me3 (purple). Values are scaled per modality (RNA, H3K4me3 and H3K27me3), mean values are shown in supplementary fig1 b. d) Uniquely mapped transcripts (x-axis) and chromatin reads (y-axis) recovered for K562 cells treated for H3K4me3 (green) or H3K27me3 (purple). Pale dots represent QC-fail cells.

We first used T-ChIC to profile the transcriptome together with either H3K27me3 or H3K4me3 distribution in K562 cells, and assess the specificity and sensitivity of the resulting signal. We observed specific enrichment of transcript reads over gene bodies, with H3K4me3 covering promoter regions of transcribed genes and H3K27me3 covering non-transcribed regions (Figure 1B, S1A). To quantify the co-occurrence of H3K4me3 and H3K27me3 signal with transcription, we grouped genes into quantiles according to their expression and determined the average coverage of both HPTMs per cell in each quantile. While highly expressed genes show co-occurrence with H3K4me3, lowly or non-expressed genes show a higher H3K27me3 coverage, which corresponds with the known active and repressive nature of these modifications (Figure 1C, S1B). To verify that both modalities do not interfere with each other, we compared signal distributions of the combined protocol with their corresponding individual method. Indeed, transcript recovered with T-ChIC correlated well with VASA-seq results, and chromatin modification distributions profiled with T-ChIC correspond to results obtained with sortChIC (Figure S1C). Additionally, the recovered number of transcript reads was independent of the number of ChIC reads and type of HPTM recovered, excluding significant interference between the two modalities (Figure 1D). Finally, we compared our multi-omic method with recently published protocols profiling both transcriptome and chromatin modifications in single cells ^29,30,32^. Both in terms of RNA and ChIC signal, our protocol recovers the highest number of fragments (Figure S1D). In addition, although all methods showed similar enrichment of H3K4me3 on active genes, T-ChIC was the only method recovering the expected anti-correlation between the repressive H3K27me3 and gene transcription without the need to correct for non-specific background signal (Figure S1E).

### Application of T-ChIC to an *in vitro* system for embryonic development recovers detailed cell types from the three germ layers

Next, we applied T-ChIC to gastruloids, assemblies of mouse embryonic stem cells (mESCs) that over the course of five days grow out into polarised structures with an anteroposterior axis and cell types derived from all three germ layers ^13,16^. We chose this system because of its scalability and faithful simulation of various processes occurring during *in vivo* embryonic development, including differentiation of cells into a complex mixture of cell types derived from all three germ layers. To follow the dynamics of HPTMs during prolonged differentiation, we optimised culture conditions and extended the protocol until 168 h after aggregation (AA) with consistent morphologies of resulting gastruloids ^33^. We achieved this by embedding the gastruloids in 10% matrigel at 96 h AA as reported before ^19,20^, with optimised embedding and media changes to improve scalability (see methods). Using this updated protocol, we generated gastruloids and performed T-ChIC experiments across a time course from 72 until 168 h AA with 24 h intervals (Figure 2A). We also included mESCs at the time point of aggregation as a starting point. We focused on H3K4me3 and H3K27me3 in this work, because of their central role in developmental gene regulation ^34,35^ and their essential role during embryonic development ^5^.

**Figure 2.**
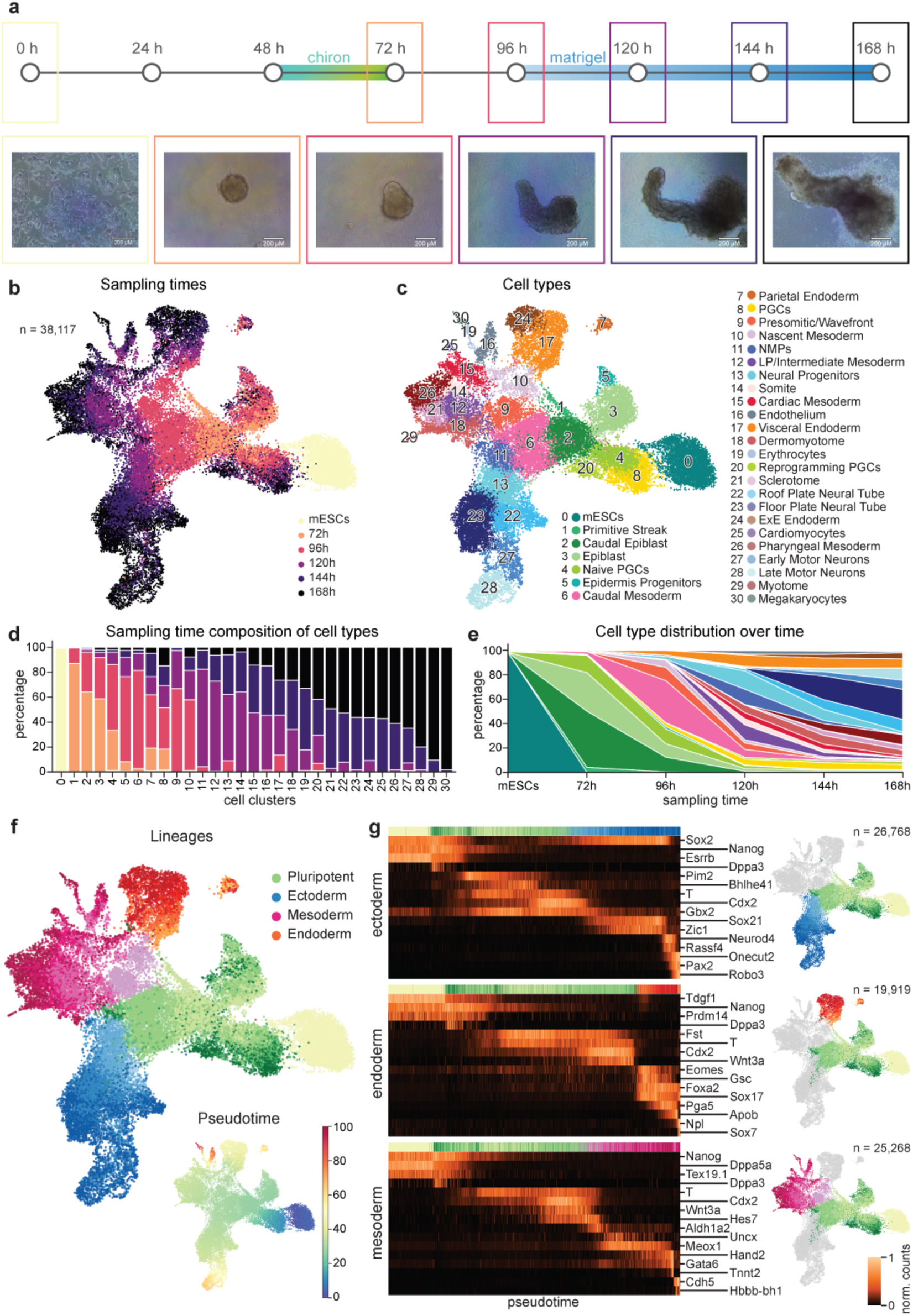
Time course of gastruloid development recovers lineage commitment to the three germ layers. a) Overview of sampling points during the gastruloid time course and the representative gastruloid morphologies. b) Transcriptome UMAP of the gastruloid dataset (n = 38,117) coloured by sampling time point of cells profiled. c) Transcriptome UMAP coloured by identified cell types. Pluripotent cell types are shaded in green, endoderm cell types are shaded in orange, mesoderm cell types are shaded in red/purple, and ectoderm cell types are shaded in blue. d) Sampling time point distribution per cell type. Numbers on the x-axis correspond to numbered cell types in c. e) Cell type distribution across time points sampled. Colours represent cell types in c. f) Transcriptome UMAP coloured by lineage. Insert: transcriptome UMAP coloured by pseudotime. g) Gene expression patterns across pseudotime for marker genes of ectoderm (top, n = 26,768), endoderm (middle, n = 19,919), and mesoderm (bottom, n = 25,268). All trajectories include pluripotent cells.

First, we confirmed that the transcript data quality was equally high for all profiled time points and HPTMs. We recovered similar numbers of total counts as well as genes detected, and did not observe major differences in the percentage of mitochondrial reads per cell (Figure S2A). The generated transcriptome UMAP consists of 38,117 single cells, with 18,279 H3K4me3-profiled cells and 19,838 H3K27me3-profiled cells intermingled in UMAP-space (Figure S2B). We observed clear transitions from early to late time points with three major trajectories and a distinct cluster representing mESCs (Figure 2B). We performed detailed cell typing based on expression of known marker genes and identified the expected diversity of cell types derived from the three germ layers, including the previously reported cardiac mesoderm, somites, endoderm, PGC-like cells and neural progenitors (Figures 2C-E, S2C-D) ^13,19,20^. Notably, especially in the later time points we recovered previously unrecorded cell types including erythrocytes and motor neurons, as well as several neural tube subtypes, emphasizing the potential of the extended culture protocol. Broadly, the identified trajectories correspond to the three germ layers - endoderm, mesoderm, ectoderm (Figure 2F). As expected, early time points mostly contribute to pluripotent cells, and after 120 h AA the recovered cells contribute to the three germ layers (Figure S2E, 2D-E). To further characterise these differentiation trajectories into more fine-grained steps than the 24 h intervals of experimental sampling, we calculated pseudotime and characterised gene expression patterns of pluripotency and differentiation markers of the three lineages (Figures 2F-G). Indeed, for all three lineages we see expression of pluripotency markers early in pseudotime, followed by gene expression of symmetry breaking factors Brachyury (*T*) ^36^ and *Cdx2* ^13,37^, and finally sequential activation of lineage specifying genes such as *Neurod4* (ectoderm), *Sox17* (endoderm) ^38^, and *Meox1* (mesoderm) ^39,40^. Lastly, we confirmed that for all identified lineages, similar ratios of H3K4me3- and H3K27me3-profiled cells are recovered (Figure S2F).

### T-ChIC recovers epigenetic regulation during gastruloid development

Next, we investigated how both recovered chromatin modalities H3K4me3 and H3K27me3 change over gastruloid development. We chose these HPTMs because they exhibit opposing functions and are known for their functional relevance and dynamic regulation during embryonic development ^5,34,35,41–45^. For both modifications, we aggregated counts across the whole genome in 50 kb windows, performed Latent Semantic Analysis (LSA) and UMAP clustering and compared obtained results to the transcriptome-based cell classification (Figure 3A). We recover similar groups for both H3K4me3 and H3K27me3, separating the three germ layers and pluripotency clusters. Notably, H3K27me3 shows a clear split between pluripotency cell types and the three lineages, while the H3K4me3 signal mainly separates mESCs and PGC-like cells and shows a more continuous transition from pluripotency to the differentiated cell types (Figure 3A). In general, we recover more H3K4me3 fragments in mESCs, while all other cell types display comparable amounts over the experimental time course. In contrast, the H3K27me3 signal continuously decreases over differentiation (Figure S3A). To select cells with high-quality chromatin profiles, we removed cells with a low total read count. Next, we identified commonly enriched bins in 200 transcriptome-based nearest neighbors and calculated per single cell the fraction of ChIC reads in these commonly enriched bins. For both modalities, ∼80% of single cells pass our thresholds (Figure S3B). Coverage plots verify our cell selection method, since selected cells show clear local enrichments, while removed cells display aspecific patterns with low enrichment (Figure S3C).

**Figure 3.**
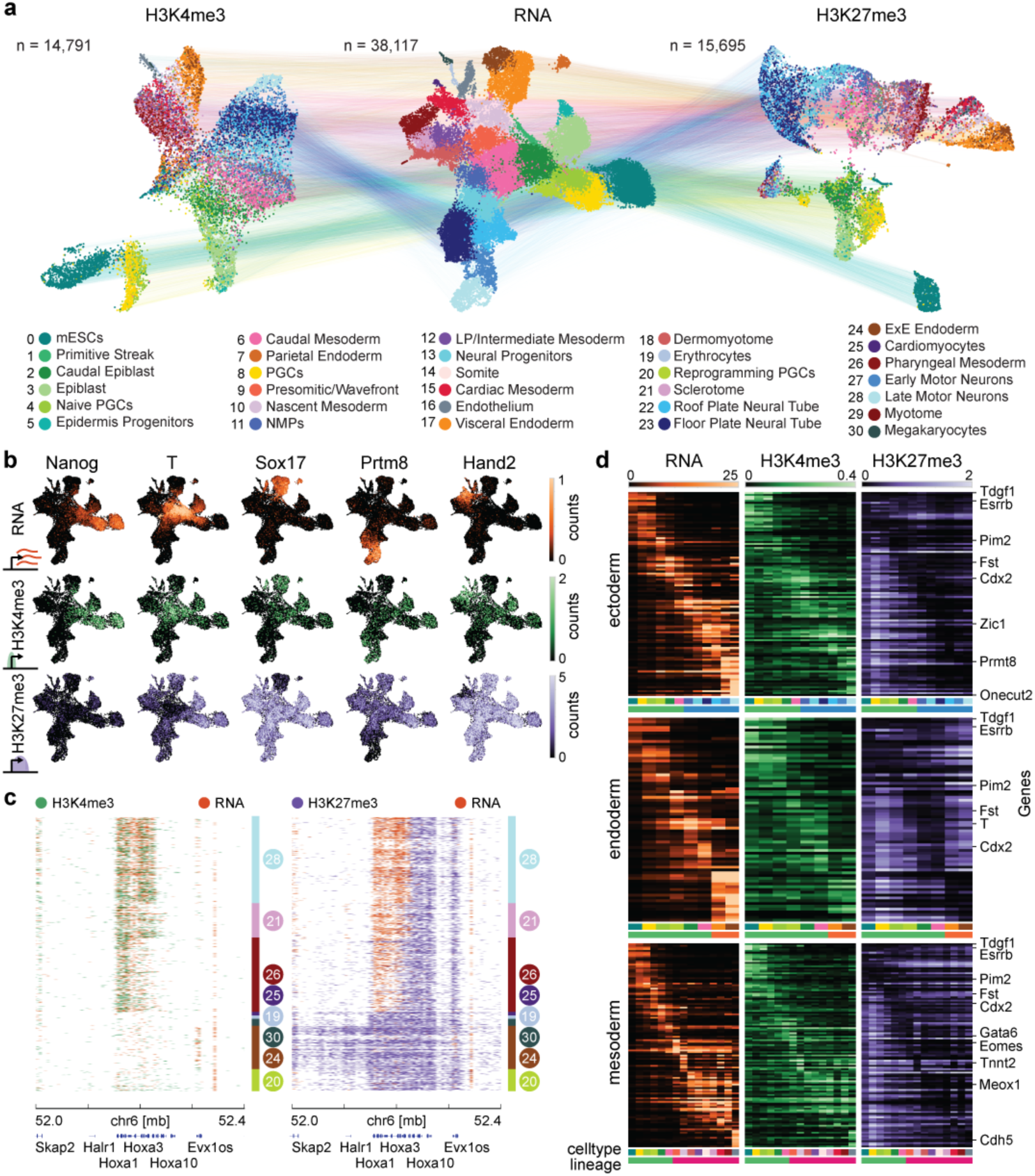
T-ChIC recovers epigenetic changes underlying transcriptional programs during gastrulation. a) Three UMAPs generated based on each of the 3 modalities. Gene expression counts are used for RNA (middle), and 50 kb genomic bins were used for H3K4me3 (left) and H3K27me3 (right). Dots are coloured based on transcript-based cell types identification (Figure 2c). b) Transcript-based UMAPs coloured by gene expression counts for RNA, promoter counts for H3K4me3 (±2.5 kb around the TSS) and gene counts for H3K27me3 (10 kb inside the gene) of selected marker genes. c) Single-cell coverage tracks of the HoxA cluster on chromosome 6. Stacked colour bar (right) indicates the relative abundance of the selected cell clusters from a. Left: RNA (orange) and H3K4me3 (green), right: RNA (orange) and H3K27me3 (purple) are shown. d) Heatmaps of mean RNA counts (orange), H3K4me3 promoter counts (green) and H3K27me3 gene counts (purple) are shown over top 5 differentially expressed genes per cell type along the three germ layer trajectories.

For further analysis of chromatin changes relative to genic transcription we selected 5 kb counting windows surrounding the transcription start sites (TSS) for H3K4me3 and the first 10 kb per gene downstream of the TSS for H3K27me3 (Figure S3D). Visualising gene-specific counts across the three modalities generally shows a co-occurrence of H3K4me3 with gene expression and anti-correlation with H3K27me3 in single cells (Figure 3B). However, while known lineage-specific genes such as *Sox17*, *Prtm8*, and *Hand2* show a strong H3K27me3 enrichment in cells not expressing the genes, the pluripotency gene *Nanog* does not show H3K27me3-mediated silencing in differentiated cell types. At the same time, H3K4me3 enrichment is not limited to cells expressing the gene but can also be found in less differentiated cells of the same trajectory (Figure 3B).

One of the most prominent examples of epigenetic gene regulation in development is the collinear derepression of the Hox gene cluster, which was also described in gastruloids in a bulk time course experiment ^13^. In single cells from 168 h AA gastruloids, we observed cell type-specific derepression of the HoxA cluster as shown by relative enrichment of RNA and HPTM distributions (Figure 3C, S3E). Pluripotent and endoderm cells are characterised by the absence of transcript or H3K4me3, combined with H3K27me3 coverage over the full length of the HoxA cluster. Conversely, mesoderm and ectoderm cells show a stepwise reduction of the H3K27me3 signal accompanied by the activation of the underlying Hox genes. To further expand this analysis, we identified the top 5 differentially expressed genes per cell type and tracked each modality along the three differentiation trajectories (Figure 3D). This revealed that most genes that are expressed during late time points are repressed by H3K27me3 in pluripotent cells. In contrast, genes expressed in pluripotent cell types accumulated H3K27me3 in a germ layer-specific manner, with endoderm cells silencing most pluripotency genes, mesoderm cells silencing a subset, and ectoderm cells accumulating repressive H3K27me3 only minimally. This observation is consistent with the genome-wide abundance of the three modalities per cell type (Figure S3F). Here, we find only moderate differences for the transcriptome and H3K4me3 signal, but large variation in H3K27me3 abundance. Pluripotency and endoderm cell types show the highest accumulation of H3K27me3, mesoderm cell types display intermediate H3K27me3 levels and ectoderm cell types show the lowest H3K27me3 counts.

### Global chromatin changes and a switch in transcription start site (TSS) usage mark the exit from pluripotency

Considering the large changes in overall H3K27me3 abundance, we decided to analyse the dynamic nature of all modalities across the trajectories in more detail. While both modalities decrease in abundance upon the start of differentiation, the decrease of H3K27me3 is more pronounced (Figure 4A, S4A). In addition, the abundance of H3K27me3 increases in endoderm specifically later in pseudotime (Figure 4A, S4A). To analyse the co-occurrence of a HPTM with RNA production of the same gene, we used a mixture model to binarise the coverage 2.5 kb upstream and 7.5 kb downstream of each TSS across cell type-specific pseudobulks (Figures 4B, S4B-C). Since our method reliably profiles the whole length of the RNA, we use only unspliced and therefore nascent transcripts for this analysis to accurately capture current transcriptional activity. The percentage of genes positive for each modality in the defined pseudobulks confirms earlier genome-wide observations of abundance per modality (Figure S4C). While the number of transcribed genes is stable across our dataset, mESC pseudobulks contain a modest increase in H3K4me3, and H3K27me3 displays marked cell type-specific differences with its abundance ranging from 70% in mESCs to 10% in ectoderm (Figure S4C).

**Figure 4.**
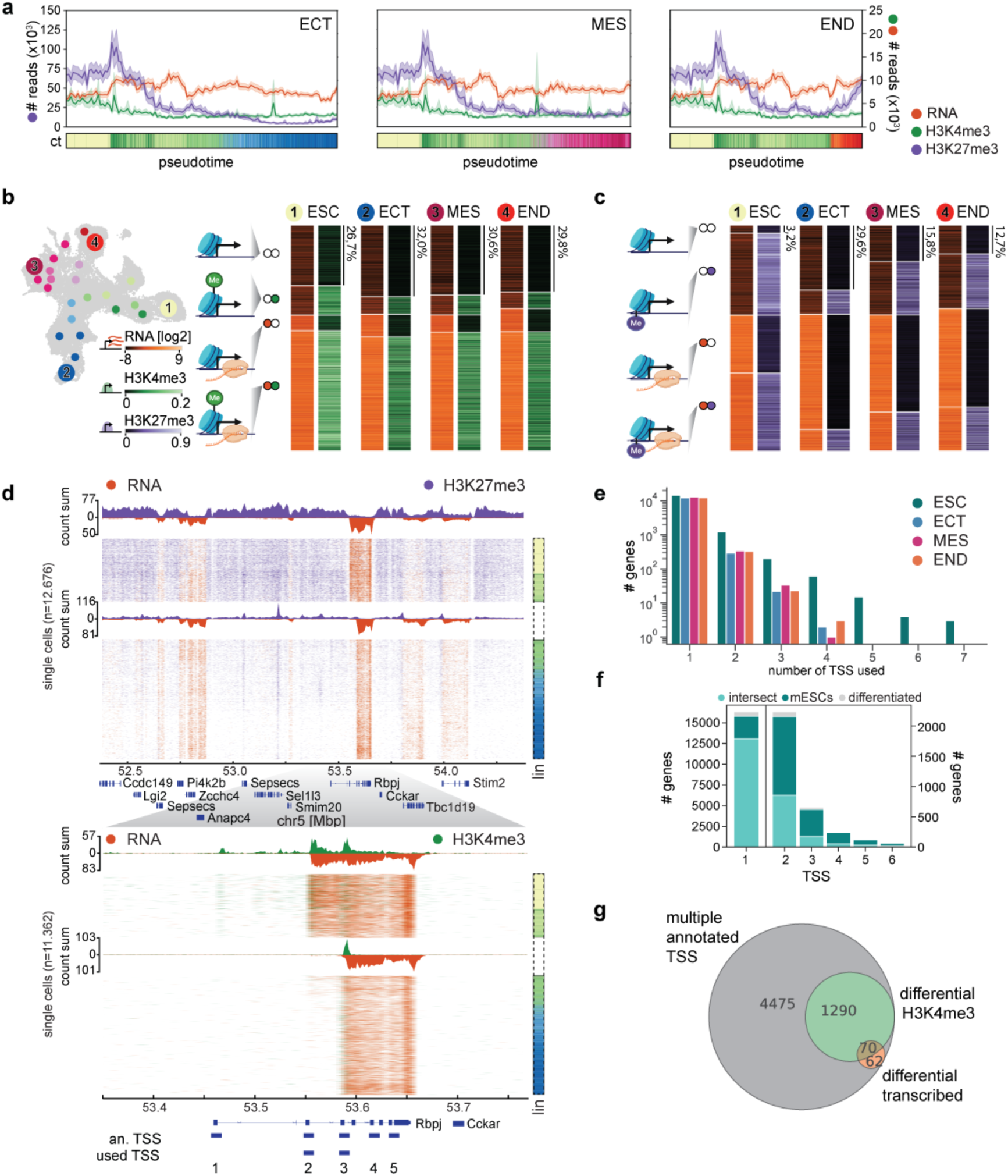
The exit of pluripotency is characterised by a global decrease in both chromatin marks and a switch in transcription start site (TSS) usage. a) Total fragments recovered per cell for RNA, H3K4me3 and H3K27me3 over pseudotime across the three lineages. b-c) UMAP depicting all pseudobulks (left) and corresponding gene states for H3K4me3 (b) or H3K27me3 (c) with RNA for selected pseudobulks. A graphic of the gene state (left) and a heatmap of total counts per gene and per state is shown. d) Single-cell coverage tracks of RNA (orange) and H3K27me3 (purple) across a 2 mb region on chromosome 5 (top), and RNA (orange) and H3K4me3 (green) across a 0.4 mb region on chromosome 5 (bottom). Cells are ordered along pseudotime across the ectoderm lineage in ascending order. Right bar: lineage (lin), including mESCs (yellow), pluripotency (green) and ectoderm (blue) cells, shaded by day. e) Bar graph showing the number of TSS used per gene in mESCs (green), ectoderm (blue), mesoderm (pink), and endoderm (orange), defined by H3K4me3 coverage. f) Stacked bar graph indicating the number of TSS per gene marked by H3K4me3 in mESCs, differentiated cells of any of the three lineages, or both (intersect). g) Venn diagram showing the overlap between genes with multiple annotated TSS, genes that differentially mark multiple of these promoters and genes where transcription initiates at different TSS between germ layers.

Next, we classified gene states by combining HPTM and transcript presence. In mESCs, the fraction of H3K4-methylated non-expressed genes is slightly larger compared to differentiated pseudobulks, while the fraction of genes covered by neither RNA nor H3K4me3 increases from mESCs to differentiated cells (Figure 4B). For H3K27me3, we observe that mESCs contain a larger fraction of genes with co-occurring H3K27me3 and RNA and H3K27me3 only, whereas differentiated pseudobulks contain a larger fraction of unmarked genes and genes with RNA but no H3K27me3 (Figure 4C). When comparing the gene states classified for both HPTMs, we observe more pronounced germ layer-specific differences in H3K27me3 states compared to H3K4me3 states (Figures 4B-C, S4D-E) The gene state with co-occurring H3K27me3 and RNA seems to be functionally relevant as H3K27-methylated genes show lower expression compared to unmethylated genes (Figures S4D-E).

To understand where the difference between HPTM abundance and number of marked genes is originating from, we expanded our analysis outside TSS regions. While H3K27me3 displays a gradual decrease across the whole genome over differentiation, H3K4me3 signal is mainly recovered surrounding genes (Figure 4D). To assess genome-wide changes in HPTM abundance we plotted Lorenz curves, which show the cumulative distribution of signal relative to genome coverage (Figure S4F). Beyond the HPTM-specific difference in genome-wide coverage, we observed only minimal changes in genome-wide enrichment for H3K4me3 between mESCs and the three lineages, while H3K27me3 showed lineage-specific differences in signal enrichment, consistent with cell type-specific differences in H3K27me3 abundance observed before (Figure S3F). When looking at an example gene, Rbpj, we observed that while H3K4me3 is recovered in coding regions, pluripotent cells display multiple H3K4me3-enriched sites per gene while differentiated cells display only one enrichment peak (Figure 4D). These additional H3K4me3 domains co-localise with annotated alternative TSS and correspond to a change in Rbpj RNA isoforms (Figure 4D), displaying H3K4me3 enrichment and transcription at TSS 2 and 3 in mESCs, followed by a switch to only TSS 3 in differentiated lineages (Figure 4D bottom panel). We hypothesized that such a decrease of marked TSS per gene could be more widely spread across genes and could in part explain the increased levels of H3K4me3 in mESCs. To investigate this, we expanded our analysis to include all alternative TSS for all genes and classified them into marked and unmarked using a mixture model on the H3K4me3 log_2_ counts (Figure S4G).

This quantification shows a dramatic decrease in the number of genes with multiple TSS covered by H3K4me3, with mESCs containing genes with up to 7 covered TSS and differentiated clusters mainly displaying one marked TSS per gene (Figure 4E). This approach identified 4.531 mESC-specific H3K4-methylated TSS, compared to only 649 TSS specific to differentiated cells and 14.195 shared TSS (Figure 4F). To understand more about the functional relevance of these extra H3K4me3 enrichments on transcription initiation, we identified genes with lineage-specific H3K4me3 enrichment and corresponding transcription initiation on alternative TSS. To identify lineage-specific H3K4me3 enrichment at the TSS that cannot be explained by the gene’s overall H3K4 methylation, we used a generalised linear model (GLM). For RNA we used a similar approach, analysing RNA reads mapping to a 10 kb window downstream of every annotated TSS and identifying genes where lineage-specific TSS changes cannot be explained by the total expression of the gene. This identified 1.360 genes with differential TSS H3K4me3, while only a subset of genes (n = 70) also exhibits transcriptional TSS switching (Figures 4G, S4G-H). This is exemplified by Bend3, Nav2 and Robo1, which show H3K4me3 coverage across multiple TSS during pluripotency but switch TSS in differentiated cells with Nac2 and Robo1 undergoing a corresponding change in RNA recovery (Figure S4H). Most genes with multiple annotated TSS only initiate transcription from one TSS (n = 3.520), as defined by the co-occurrence of both H3K4me3 enrichment and RNA starting from the same TSS, with similar abundance of RNA along the gene in different germ layers (Figure S4I). However, for 411 of these genes, the main annotated TSS is not used in our gastruloid system and an alternative TSS is active instead (Figure S4J-K). To ensure correct determination of the TSS chromatin state in downstream analyses, we used a corrected annotation based on our experimental data in the rest of the manuscript.

### Classifying genes into discrete states uncovers differential epigenetic gene regulation in the three germ layers during differentiation

The dynamic changes in both H3K27me3 and H3K4me3-marked TSS, contrasting a stable fraction of transcribed genes, led us to interrogate the interplay between active and repressive chromatin states. For this, we combined the binary classification of all three modalities into 8 gene states (Figure 5A). In agreement with previous studies ^46^, the gene states with the smallest number of genes contain H3K4me3 only, transcription only, and H3K27me3 with transcription (Figures 5B-C, S5A-B). The most abundant state (∼40% of all genes) in all cell types is the combination of transcription with H3K4me3, which reflects actively transcribed genes such as housekeeping genes. We also define the bivalent state, which is marked by co-occurrence of active H3K4me3 with repressive H3K27me3 and is thought to prime lineage-specific genes in pluripotent cells for activation later during development ^41,42,44,47^.Interestingly, the majority of bivalent genes are lowly transcribed (Figure S5A).

**Figure 5.**
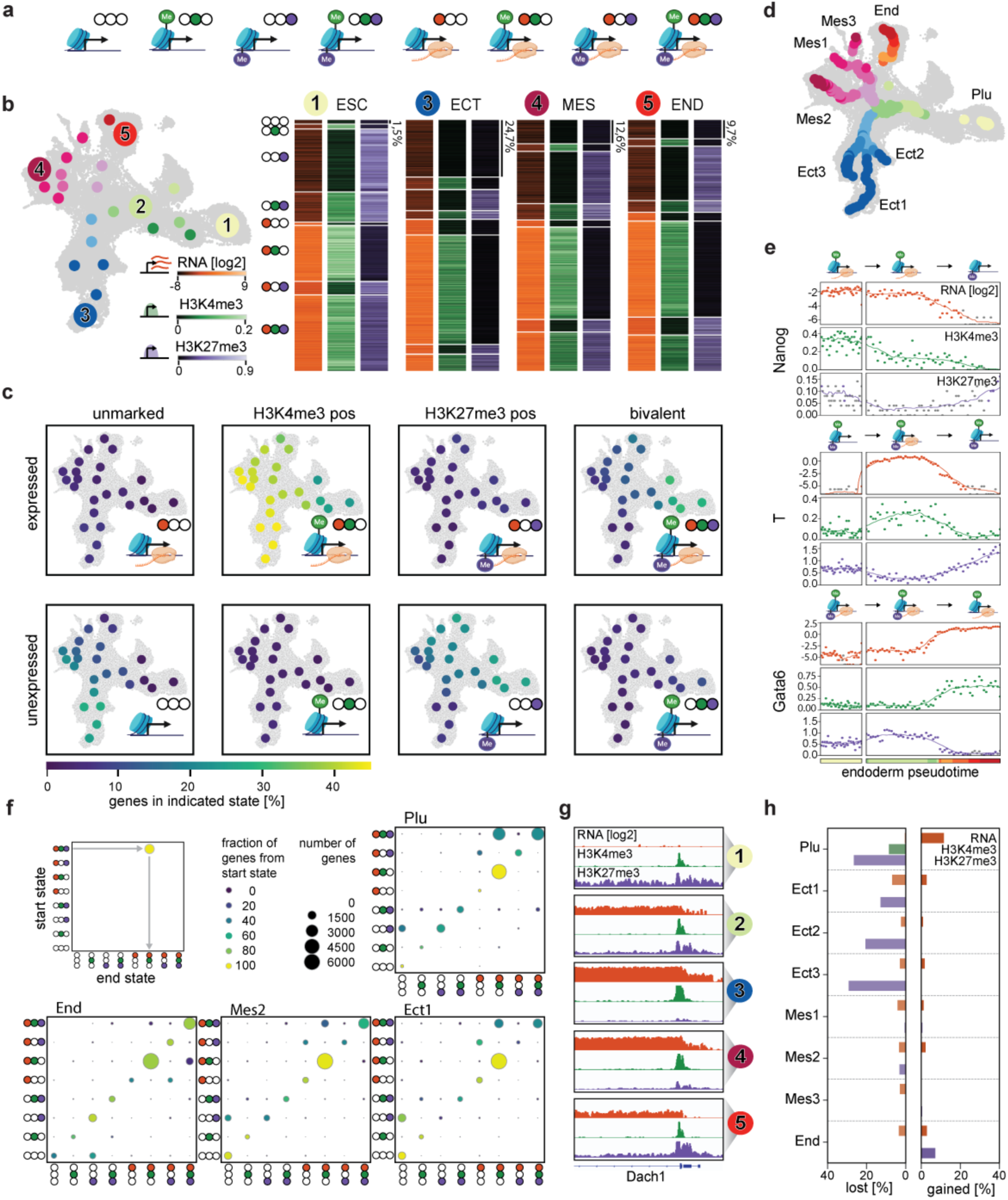
mESC-specific chromatin states resolve in a germ layer-specific manner. a) Graphical representation of 8 gene states, representing the possible combinations of the three recovered modalities. b) left: UMAP showing cell type-specific pseudobulks used for gene state analysis. Numbers refer to heat maps (right) and coverage tracks shown in g. Right: Heatmaps of the four extremities of the UMAP showing log_2_ gene expression (left, orange), promoter H3K4me3 (middle, green) and H3K27me3 (right, purple). Rows indicate individual genes ordered by gene state. c) UMAPs with pseudobulk dots representing the centroid of each cluster coloured by the fraction of genes in the indicated gene state. d) Detailed pseudobulk trajectories for the three germ layers and pluripotency. e) State changes over pseudotime for example genes Nanog, T, and Gata6, along endoderm trajectory. Dots indicate pseudobulks, colours represent whether the representative modality is classified as present (orange, RNA; green, H3K4me3; purple, H3K27me3) or absent (grey). f) Summary of state changes for the pluripotent trajectory and one example trajectory per germ layer. Dot size indicates number of genes per state change, colour indicates fraction of genes with the same start state changing to indicated state. g) Pseudobulk tracks of example gene Dach1 across three indicated pseudobulks along ectoderm differentiation (see a). h) Number of genes with significant increase (‘up’) or decrease (‘down’) of indicated modality (orange, RNA; green, H3K4me3; purple, H3K27me3) for each of the 8 trajectories.

Four gene states are especially dynamic across cell types, including transcribed bivalent genes (Figures 5C, S5A). The number of genes for which we recover this state is strongly enriched in pluripotent cells, with a lower abundance in later time points (Figure 5C). This distribution is mirrored in the higher abundance of the H3K4me3-positive expressed gene state in differentiated cell types. We also observe more lineage-specific enrichments of gene states, such as fully unmarked genes in the ectoderm lineage and H3K27me3-only in the pluripotent and endoderm lineages (Figure 5C). This change in the abundance of H3K27me3 containing states, is further reflected by a time point- and germ layer-specific switch in genome-wide distribution (Figures S5B-C). In mESCs and endoderm, H3K27me3 spread out across whole gene bodies. Upon quantification of H3K27me3 abundance in TSS regions compared to total abundance, pluripotent and endoderm cells indeed show a low fraction of reads falling in TSS regions (Figure S5C). In contrast, ectoderm and mesoderm display a TSS-focused distribution and a larger fraction of H3K27me3 reads in TSS regions, corresponding to a depletion of H3K27me3 in gene bodies and intergenic regions (Figure S5B-C).

The dynamic changes in gene states and genomic distribution of HPTMs are in line with previous studies describing a decrease of H3K27me3 abundance during mESC differentiation using mass spectrometry and quantitative ChIP ^48,49^. Similarly, in vivo studies of whole mouse embryos showed that at the blastocyst stage, embryos undergo widespread epigenetic reprogramming including the removal of parental epigenetic markings, loss of DNA methylation and a temporal spread of polycomb-mediated H3K27me3 and H2AK119Ub decoration ^50^.

We reasoned that such global changes in HPTM distribution should be reflected by the composition of the involved writer and eraser complexes. We therefore compared the expression of known epigenetic regulators of H3K4me3 and H3K27me3 in the different lineages as well as corresponding de-methylases (Figure S5D). Upon differentiation of mESCs, we find a change in the relative abundance of half of the COMPASS components, while corresponding de-methylases appear unchanged. In contrast, Polycomb Repressive Complex 2 (PRC2) group components display various changes: mESCs highly express PRC targeting components Eed and Jarid2, pluripotent and endoderm cells highly express Rbbp7, and mesoderm and ectoderm cells have relatively low expression of most PRC2 components with the exception of Rbbp4 and the enzymatic core Ezh2. In agreement with the observed global reduction of H3K27me3, we also observe increased levels of its demethylase Kdm6b in differentiated cells. Together, the lineage-specific changes in writer and eraser components mirror the observed chromatin state usage, supporting a systematic change in epigenetic regulation at this stage of embryonic development.

To study how the observed epigenetic states are connected during gene regulation, we set out to define gene state changes. First, we defined more detailed trajectories to reflect the full biological and temporal complexity of the underlying data (Figures 5D, S5E). Known dynamic marker genes, such as Nanog, T, and Gata6 ^51^ display the expected behaviour across these trajectories, which is reflected by a switch in gene state (Figure 5E). Next, we summarised all gene state changes across all trajectories (Figure 5F). In all trajectories, we recover stably expressed housekeeping genes as the biggest fraction, starting and ending as H3K4me3-covered and expressed. Further, the pluripotency trajectory is dominated by a loss of H3K27me3, with genes changing from bivalent expressed to active, and from H3K27me3-covered to fully unmarked (Figure 5F). This trend is continued along the ectoderm trajectory, which is characterised by a loss of repressive H3K27me3, often without a corresponding change in transcription, and is exemplified at the single-gene level (Figures 5F, S5E-F). The majority of bivalent genes are thereby already expressed before H3K27me3 is fully lost, exemplified by the large fraction of bivalent expressed genes (Figures 5C,F) including Gata6 and Dach1 (Figures 5E,G). In contrast, the endoderm trajectory is marked by a *de novo* gain in H3K27me3, while mesoderm trajectories display mixed behaviour, both resolving a subset of bivalent genes to fully active as well as re-gaining repressive signatures on other genes (Figures 5F).

Performing a conventional differential abundance analysis along these trajectories recovers the same global germ layer-specific trends (Figures 5H, S5G), but also highlights the dynamic changes that occur inside stable states. For example, genes losing H3K27me3 from a bivalent state in the pluripotency trajectory often display an increase in transcription strength, reflecting a de-repression of the underlying gene. In the endoderm trajectory we observe gene states with H3K27me3 to accumulate even higher levels of this mark, further consolidating the repressive state (Figure S5G, bottom panel).

### Germ layer-specific genes sequentially lose H3K27me3 over gastrulation

To understand how dynamic changes in chromatin state contribute to lineage commitment, we defined differentially expressed genes between the pluripotent cluster and each of the three germ layers using DESEQ2 and analysed their behaviour over differentiation (Figure 6A; ^52^). For each comparison, we selected significantly changing genes (p-value <0.05, log-fold change >2) exclusive to one germ layer (Figure 6B-C). Gene ontology (GO) analysis shows the expected biological terms enriched in the three gene sets, with ectoderm genes (n = 385) enriched for “neurogenesis”, mesoderm genes (n = 417) enriched for “circulatory system development”, and endoderm genes (n = 622) enriched for general developmental terms like “anatomical structure morphogenesis” (Figure S6A).

**Figure 6.**
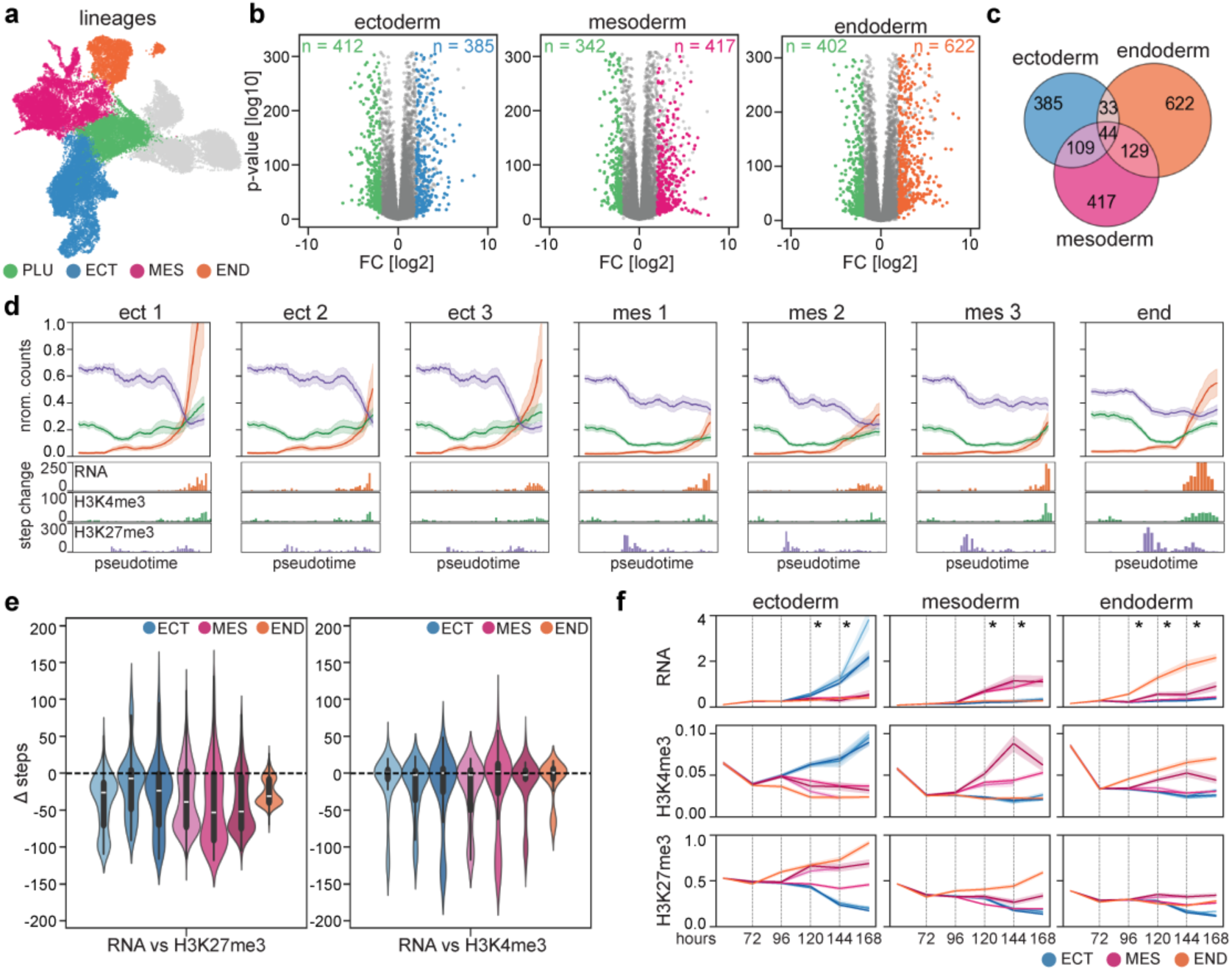
Sequential de-repression of bivalent genes underlies lineage-specific gene activation. a) UMAP indicating the cell populations used for differential gene expression analysis with DEseq2. b) Volcano plots displaying significant differentially expressed genes recovered for each germ layer. c) Venn diagram displaying the overlap between differentially expressed gene sets of the three lineages. In subsequent analysis, only differentially expressed genes unique for each lineage were used (ectoderm: n = 385, mesoderm: n = 417, endoderm: n = 622). d) Top: line plots indicating abundance of RNA (orange), H3K4me3 (green), and H3K27me3 (purple) of differentially expressed gene sets over pseudotime across the 7 trajectories. Bottom: histogram indicating the step at which genes of the corresponding gene set undergo a significant change (2x std) for each modality (orange, RNA; green, H3K4me3; purple, H3K27me3). e) Violin plots of the temporal delay between significant chromatin relative to transcription changes. For RNA and H3K4me3 the time point of significant increase was determined. For H3K27me3 the time of decrease was used. f) Line plots indicating abundance of RNA (top), H3K4me3 (middle), and H3K27me3 (bottom) of differentially expressed gene sets over sampling time (left: ectoderm-specific genes, middle: mesoderm-specific genes, right: endoderm-specific genes). The 7 trajectories are indicated as lines: ectoderm: light to dark blue, mesoderm: light to dark pink, endoderm: orange.

For each of the differentially expressed gene sets, we analysed their gene state transitions along the 7 differentiation trajectories (Figure S6B). Differentially expressed genes generally start in a bivalent state in mESCs and either resolve to a fully expressed state (in ectoderm, Figure S6B left panel), or retain H3K27me3 despite transcriptional activation (in endoderm, Figure S6B right panel). This mirrors the general behaviour observed for all genes, highlighting de-repression of bivalent genes as the major form of gene activation during gastruloid development.

To obtain more detailed information about the order of change between the different modalities, we decided to look beyond absolute states into more dynamic differences between HPTMs and RNA. We visualised the average abundance of all three modalities on differentially expressed genes over pseudotime in the lineage of their activation (Figure 6D). As expected from the state transitions, ectoderm genes generally lose more H3K27me3 and gain higher levels of transcription over pseudotime, whereas endoderm and mesoderm genes retain some H3K27me3 even after transcription activation occurred. To identify the time point of significant change per modality, we determined per gene the first pseudotime step in the trajectory where we observe a change beyond two standard deviations from the mean (Figure S6C). For RNA and H3K4me3 we determined the first point of significant increase, while for the repressive H3K27me3 we determined the point of significant decrease (Figure S6D). To identify whether the modalities change at different steps in pseudotime, we subtracted the step of significant increase in RNA from the step of significant chromatin change (increase for H3K4me3 or decrease for H3K27me3; Figure 6E). For all trajectories, we observed that H3K27me3 is removed before transcription activation (Figure 6E, left panel). In addition, we observe a longer delay between H3K27me3 loss and transcriptional activation for mesoderm- and endoderm-specific genes compared to ectoderm-specific genes in their respective trajectory. This shift cannot be explained by technical limitations, such as sparsity or signal separation, as the even sparser H3K4me3 shows perfectly timed changes together with transcription (Figure 6E, right panel). This suggests that while mesoderm and endoderm-specific genes depend on other stimuli besides loss of H3K27me3-mediated repression for their transcriptional activation, ectoderm genes are primed for transcriptional activation through loss of H3K27me3 alone.

To understand how germ layer-specific differentially expressed genes are epigenetically regulated in alternative lineages, we analysed their behaviour in all seven trajectories (Figures 6A-C, F). As expected, transcription and H3K4me3 display the largest increase in the intended germ layer, while H3K4me3 is progressively lost in alternative trajectories. H3K27me3 displays differential regulation in alternate lineages. Ectoderm genes remain repressed in mesoderm and endoderm lineages, and only lose H3K27me3 abundance in the ectoderm itself. Mesoderm genes remain highly repressed in endoderm, while both mesoderm and ectoderm lose H3K27me3 abundance across this gene set. Lastly, endoderm genes show loss of H3K27me3 abundance in all lineages. These same gene sets also show a difference in their timing of activation. Endoderm-specific genes are upregulated at 96 h AA, and mesoderm- and ectoderm-specific genes at 120 h AA (Figures 6F, S6E). This temporal shift in activation can also be seen in the state transitions from mESCs to pluripotent where 15% of endoderm-specific genes change from bivalent expressed to fully active (RNA + H3K4me3, Figure S6B right panel).

Taken together, the majority of genes that are activated over differentiation are marked as bivalent in pluripotent cells. Upon differentiation, each lineage activates its specific gene set, which further gains H3K4me3 abundance and transcription. Endoderm cells lose H3K27me3 only on genes later activated in this germ layer. Mesoderm gene-expressing cells show the additional loss of H3K27me3 on their geneset, and display the same temporal disconnect between de-repression and transcriptional activation. Finally, cells expressing the ectoderm-specific genes lose H3K27me3 on all 3 genesets, followed by immediate transcription activation of their genes (Figure 7).

**Figure 7.**
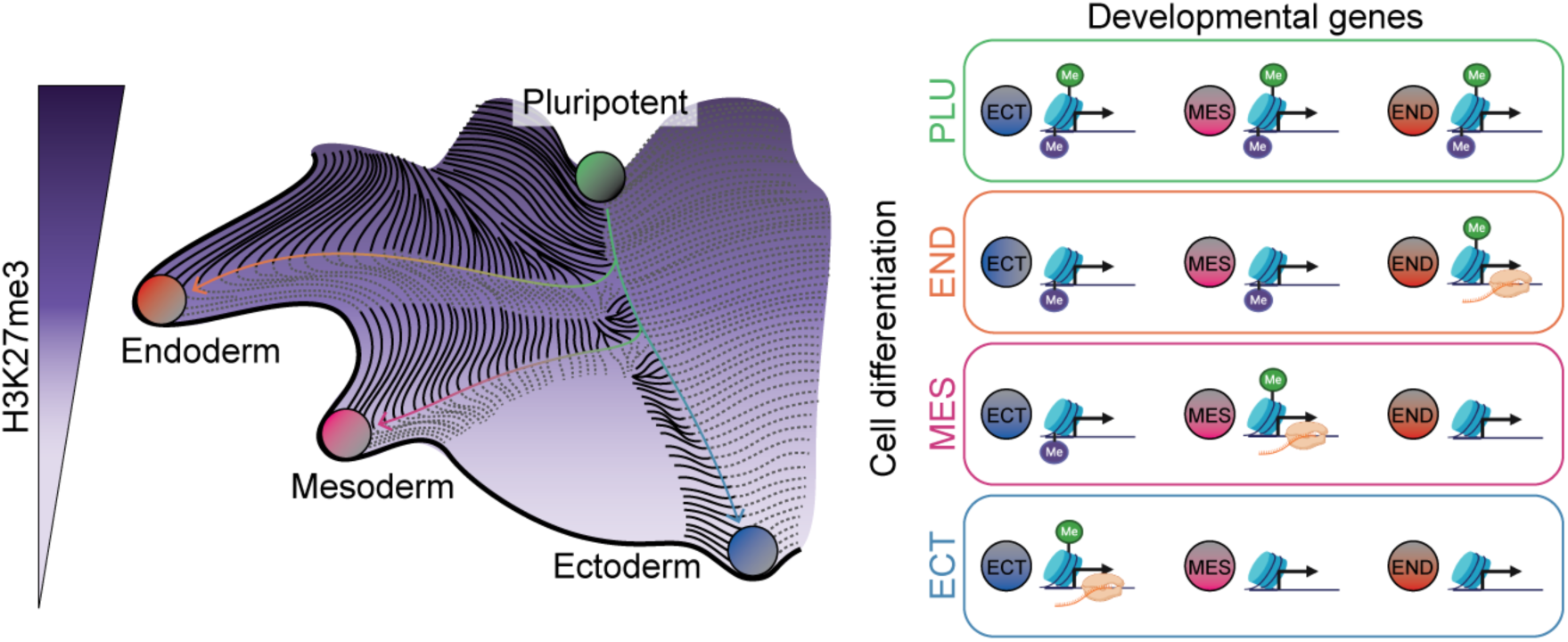
Graphical model of the sequential de-repression of lineage-specific genes. Left: Graphical depiction of the epigenetic landscape. Right: Schematic depiction of the presence of the three modalities (RNA, H3K4me3, H3K27me3) on lineage-specific gene sets in pluripotent and differentiated cells. From pluripotent to differentiated states, H3K27me3 is gradually lost. Endoderm retains H3K27me3 abundance on all but endoderm-specific genes, where this HPTM is lost early in differentiation. Mesoderm loses H3K27me3 additionally on mesoderm genes, but maintains repression of ectoderm-specific genes. Finally, Ectoderm displays the most dramatic loss of H3K27me3 abundance, showing loss of H3K27me3 on all germ layer-specific gene sets.

## Discussion

In summary, we introduce T-ChIC, a novel single-cell method that allows the recovery of the full-length transcriptome and HPTM distributions with high sensitivity and accuracy. We demonstrate that our technique is particularly faithful in the recovery of repressive chromatin modifications, and outperforms current Tn5-based technologies in terms of sensitivity. Repressive HPTMs are especially important to characterise, as they form a boundary for transcriptional gene activation. This is crucial to understand how epigenetic pathways restrict gene expression and contribute to lineage decisions. Additionally, pA-MN based approaches were shown to more faithfully profile weak genomic interactors, such as TFs, compared to Tn5-based technologies. Therefore, we envision that T-ChIC, currently the only MN-based RNA-chromatin multi-omics technique, will similarly prove to allow profiling of a more diverse set of genome-protein interaction targets.

While the T-ChIC method is currently limited to a mid-throughput range due to its processing in plates, it results in a 10-fold increase in sensitivity compared to current 10x-based methods. It therefore requires less cells to provide the same level of complexity, making it comparable in time investment and superior in terms of financial efficiency.

Here, we applied T-ChIC to an extended gastruloid system and profiled H3K4me3 and H3K27me3 distributions over a time course of seven days. We demonstrate that differential gene expression is accompanied by corresponding changes in epigenetic modifications. This includes the Hox gene cluster, which is known to display collinear expression in gastruloids ^13,53^. We show that H3K27me3 and H3K4me3 profiles divide the *Hox* cluster into an active and an inactive compartment, which correspond to collinear activation comparable to the *in vivo* scenario. As expected, promoters of transcribed genes are covered by H3K4me3, and genes covered by H3K27me3 remain mostly unexpressed.

The pluripotent state of mESCs displays several unique characteristics. First, we observe widespread H3K4me3 accumulation on annotated alternative TSS. While the majority of these events do not lead to a change in transcription initiation, this increase in H3K4me3 distribution does indicate a fundamental change in the H3K4 methylation machinery in concordance with RNA polymerase activity. In particular, mESCs displayed multiple marked TSS per gene, which upon differentiation becomes restricted to one main preferred transcript version. While accumulation of mESC-specific intra- and inter-genic H3K4me3 foci has been reported before at unmethylated CpG islands ^54^, why the majority of these extra H3K4me3 loci has no functional consequence on transcription initiation is still unclear.

The most striking change we observe is a genome-wide reduction of H3K27me3 levels at the onset of differentiation. While this massive loss of repressive chromatin was counter-intuitive, considering the accessible chromatin state reported in mESCs ^45,55^, bulk experiments *in vivo* and *in vitro* have found a comparable loss of H3K27me3 both at the exit of blastocyst development ^50^, and during the transition from ground state to naive pluripotency ^48,49^, respectively. The majority of H3K27me3 loss occurs independent of transcriptional activation, and potentially reflects the derepression of genes for later activation. Additionally, we observe a large fraction of developmentally associated genes that are bivalently covered in mESCs, as previously described ^41,44,56^. We identify loss of H3K27me3 on bivalent genes as the most abundant form of gene activation, including for genes expressed in a lineage-specific manner. Complete loss of H3K27me3 does not seem to be essential for gene activation, as a large fraction of genes is lowly expressed in a bivalent state. Especially in the mesoderm and endoderm, these genes remain bivalently marked but do gain additional transcriptional output. Loss of H3K27me3 on bivalent promoters on the other hand does not always lead to a direct gene activation, as we observe a delay between the loss of H3K27me3-mediated repression and transcriptional activation. This supports previous work that describes the role of bivalency mainly to allow future activation by preventing the deposition of DNA methylation ^57,58^, compared to a poised state that ensures fast activation after H3K27me3 is lost.

Based on our results, we propose a model for epigenetic perceptiveness towards cell fate determination (Figure 7). While cells are primed for ectoderm specification upon release from pluripotency ^59^, our data indicates that cells become epigenetically perceptive towards endoderm first. In our model system, we observe that endoderm-specific genes become de-repressed earliest in time as indicated by loss of H3K27me3 (at 96 h AA), but transcriptional activation of the same gene set lags behind. Next, cells become perceptive towards the mesoderm lineage, as indicated by additional loss of H3K27me3 on the mesoderm gene set and a similar delay in transcription activation. We hypothesize that while cells require the de-repression of endoderm- and mesoderm-specific gene sets, another stimulus is required for cells to initiate transcription activation, leading to lineage commitment ^14,60,61^. The activation of the Wnt pathway is a likely candidate to perform this function ^14,61^. Cells become perceptive to the ectoderm lineage last, as indicated by loss of H3K27me3 on ectoderm-specific genes.

In contrast to the other two lineages, this appears to be sufficient for immediate transcriptional activation of the same genes and commitment to the ectoderm fate. This suggests that ectoderm-specific genes could be the only genes pre-set for their activation and therefore representing a true de-repression. This is in line with previous studies *in vitro* where differentiation towards the ectoderm lineage was identified as the default option ^59,62,63^. In contrast, for cells to differentiate towards a mesoderm fate, they seem to require the continued repression of ectoderm-specific genes. Lastly, endoderm appears to require active H3K27me3-mediated repression of both alternative lineages.

To conclude, we find clear differentiation stage- and germ layer-specific usage of chromatin states for gene regulation, with mESC, endoderm and ectoderm representing extremes of each class, and mesoderm displaying intermediate characteristics between ectoderm and endoderm. The observed changes thereby most likely reflect the exit of epigenetic reprogramming and corresponding ‘locking in’ of a stable epigenetic landscape, an essential process to ensure faithful lineage-specific differentiation at the onset of organism formation.

While pluripotency and endoderm cells display H3K27me3-mediated silencing of unexpressed genes, mesoderm and especially ectoderm cells show H3K27me3 enrichment only on a subset of unexpressed genes. This raises the question whether other epigenetic regulators, such as H3K9me3 or DNA methylation, might compensate for these lineage-specific differences in H3K27me3 usage. Previous work on H3K9me3 on dissected germ layer cells similarly described higher levels of H3K9me3 in endoderm and mesoderm compared to ectoderm ^64^. DNA methylation studies profiling mouse gastrulation recovered a rapid increase in global DNA methylation at the exit of epigenetic reprogramming that could replace reducing H3K27me3 levels at the onset of lineage commitment. While genome-wide DNA methylation levels do not show differences across lineages, this might not reflect their abundance on small regulatory regions such as promoters, leaving the question how ectoderm cells epigenetically repress silent genes unanswered ^59^. These results emphasise the importance of looking at chromatin regulation in a cell type-resolved manner and display the dynamic regulation of repressive chromatin during lineage commitment. Applying single-cell multi-omics techniques to study both active and repressive chromatin states will be instrumental to build a comprehensive understanding of how different epigenetic pathways cooperate in restricting transcriptional activation and thereby govern differentiation decisions.

## Methods

### Cell culture

K562 cells (ATCC® CCL-243™) were grown in RPMI 1640 Medium GlutaMAX™, supplemented with 10 % FCS, Pen-Strep and non-essential amino acids. After harvesting, cells were washed 3 times with room temperature PBS before continuing with the cell fixation. 129S1/SvImJ / C57BL/6 mESCs ^65^ were cultured in a humidified incubator (5 % CO2, 37 ℃) on 0.1 % gelatin-coated cell culture dishes in ESLIF medium (GMEM (Gibco) containing 10 % fetal bovine serum (FBS, Gibco, #16141-079, lot 2215500RP), 1 % Sodium Pyruvate, 1 % non-essential amino acids, 1 % GlutaMAX, 1 % penicillin-streptomycin, 0.2 % β-mercaptoethanol, and 1000 units/mL mouse leukemia inhibitory factor (mLIF, ESGRO)). Cells were split every second day by washing with PBS0, dissociating with TrypLE for ∼5 min, centrifuging at 300 g for 3 min, and replating the pellet 1:5 on gelatinised plates. 2i medium was prepared as follows: 48.1 % DMEM/F12 (Gibco) and 48.1 % Neurobasal (Gibco) containing 0.5 % N-2 supplement (Gibco, #7502048), 1 % B-27 supplement (Gibco, #17504044), 1 % GlutaMAX, 1.1 % penicillin-streptomycin, 0.2 % β-mercaptoethanol, 1000 units/mL mouse leukemia inhibitory factor (mLIF, ESGRO), 3uM Chiron (CHIR99021, Tocris #4423) and 1uM PD0325901 (Sigma, #PZ0162). Before gastruloid formation, mESCs were cultured for 2 days in 2i medium followed by 2 days in ESLIF medium and aggregation. All cell lines were routinely tested and confirmed to be free of mycoplasma.

### Gastruloid formation

Gastruloids were generated as described before ^16,19^ with the following adaptations: N2B27 medium (Takara, #Y40002) was passed through a 0.22 μm filter before use and 600 cells were used for aggregation instead of 300. For all time points from 96 h AA onwards, gastruloids were embedded in 10 % matrigel (Corning, #356231, lot 0006004) in N2B27 medium as follows: 6-well plates were coated with 1 mL 10 % matrigel in N2B27 and left to solidify at 37℃ for >1 h. Gastruloids were harvested at 96 h AA from 96-well plates, pooled, and placed on ice as shortly as possible. All medium was removed from gastruloid pools and replaced with 10 % matrigel in N2B27, and a maximum of 32 individual gastruloids in matrigel were plated per 6-well and distributed evenly. At 120 h AA and 144 h AA, gastruloids were either harvested or medium was refreshed by removing and adding 1 mL N2B27.

### Dissociation and fixation of mESCs and gastruloids

mESCs were dissociated as described above, resuspended in PBS0 supplemented with 5 % FBS, and filtered through a 35 μm filter (Falcon, 352235).

Non-matrigel-embedded gastruloids were pooled in protein low-bind tubes (#0030122216), washed with PBS0, and incubated with 1 mL trypsin (Gibco) for ∼5 min at 37 ℃. Gastruloids were mechanically broken up into a single-cell suspension by pipetting with a P200 pipette, resuspended in N2B27 medium, and washed twice with PBS0. Embedded gastruloids were removed from matrigel as follows: N2B27 medium was removed from 6-well plates, replaced with 1 mL Corning Cell Recovery Solution (#354253), and incubated on ice for ∼15 min. Gastruloids were flushed from matrigel solution using a P1000 pipette and collected in protein low-bind tubes, followed by dissociation as described above.

From here, all steps were performed at 4 °C. Cells were washed once in cold PBS0, resuspended at 1 million cells per 300 µL PBS0, pre-cooled ethanol was added while vortexing to a final concentration of 70 %, and cells were mildly fixed at −20 °C for 1 h. After, cells were washed once in antibody incubation buffer (47.5 mL H2O RNAse free, 1 mL 1 M HEPES with pH 7.5 (Invitrogen), 1.5 mL 5 M NaCl, 3.6 mL pure spermidine solution (Sigma Aldrich), 0.05 % Tween-20, EDTA free protease inhibitor, 1 U/µL RNasin® Plus Ribonuclease Inhibitor, 2 mM EDTA; buffer identical to T-ChIC wash buffer + 2 mM EDTA) at 4 °C, resuspended in antibody incubation buffer supplemented with 10 % DMSO, and aliquoted at 0.5 million cells per tube for long-term storage at −80 °C.

### T-ChIC protocol

#### pA-MN targeting

After thawing, cells were pelleted at 500 g for 4 min and washed once in T-ChIC wash buffer (47.5 mL H2O RNAse free, 1 mL 1 M HEPES with pH 7.5 (Invitrogen), 1.5 mL 5 M NaCl, 3.6 mL pure spermidine solution (Sigma Aldrich), 0.05 % Tween-20, EDTA free protease inhibitor, 0.2 U/µL RNasin® Plus Ribonuclease Inhibitor; buffer identical to basic T-ChIC buffer + protease and RNAse inhibitor). (2) Cells were resuspended in 200 µL antibody incubation buffer per 0.5 million cells and were aliquoted into 0.5 mL protein low binding tubes containing the primary HPTM antibody (1:200 dilution for H3K4me3 and 1:100 for H3K27me3) diluted in 200 µL antibody incubation buffer. (3) Cells were incubated overnight at 4 °C on a roller, (4) followed by washing once with 500 µL T-ChIC wash buffer. (5) Afterwards, cells were resuspended in 500 µL T-ChIC wash buffer containing a total of 1 U/µL RNAsin plus, PaMN (3 ng/mL) and Hoechst 34580 (5 mg/mL) and (6) incubated for 1 h at 4 °C on a roller. (7) Finally, cells were washed an additional 2 times with 500 µL T-ChIC wash buffer before passing through a 70 µm cell strainer (Corning, 431751) for sorting.

#### FACS sorting

(8) For all experiments, cells were gated for G1 cell cycle stage based on Hoechst staining and forward scatter and trigger pulse width to further remove cell doublets on an Influx FACS machine. Cells were sorted into 384-well plates containing 5 µL sterile-filtered mineral oil (Sigma Aldrich) and 50 nL RNA recovery primer (28 ng/mL, supplementary table 1) per well using index sorting, which records FACS information for every sorted well. To further increase sorting specificity of single cells, we used the following sort settings: objective: single, number of drops = 1, extra coincidence = complete empty (no signal in the previous and next drop), and phase mask = center 10/16 (cell is in the middle of the sorted drop).

For both marks we sorted 3 plates of K562 cells, 6 plates with mESCs, 6 plates with 72 h AA gastruloids and 12 plates for all later gastruloid timepoints. For H3K4me3, clone G.532.8 (Invitrogen, MA5-11199) was used at 1:400 dilution and for H3K27me3, clone C36B11 (NEB, 9733S) was used at 1:200 dilution. Data was collected using BD FACS software (v1.2.0.124).

#### Protein A-MN activation

Volumes for the following steps of the protocol were distributed using a Nanodrop II system (Innovadyme) and plates were spun for 2 min at 4 °C and 2000 g after each reagent addition. (9) 100 nL of Basic T-ChIC buffer (47.5 mL Nuclease-free water, 1 mL 1 M HEPES with pH 7.5 (Invitrogen), 1.5 mL 5 M NaCl, 3.6 mL pure spermidine solution (Sigma Aldrich), 0.05 % Tween20) containing 3 mM CaCl_2_, was added per well to induce pA-MN-mediated chromatin digestion. (10) For digestion, plates were incubated for 30 min in a PCR machine set at 4 °C (11) The reaction was stopped by adding 100 nL of the stop solution (40 mM EGTA to chelate Ca^2+^ and stop MN (Thermo, 15425795), 1.5 % NP40, 2 mg/mL temperature-labile proteinase K (NEB, P8111S), and 12.5 mM MgCl_2_ for RNA fragmentation). (12) Plates were incubated in a PCR machine for 20 min at 4 °C, followed by the release of chromatin and digestion of PaMN by proteinase K at 37 °C for 2 h and heat inactivation at 55 °C for 20 min. Afterwards, plates can be stored at −80 °C until further processing.

#### Library preparation

(13) After thawing, RNA was fragmented at 85 °C for exactly 2 min before cooling down on ice. (14) RNA fragments were repaired and poly-A-tailed through the addition of 150 nL RNA repair mix per well and incubation at 37 °C for 1 h. Per plate, the RNA repair mix contains 18.75 µL Nuclease-free water, 12 µL 0.1 M DTT, 0.9 µL 1 M MgCl_2_, 18 µL 1 M KCl, 16.8 µL 1 M Tris with pH 8, 4.5 µL 50 % PEG8000, 0.45 µL 20 mg/mL BSA, 5.25 µL 0.1 mM ATP, 6 µL RNase-OUT, 1.05 µL polyA Polymerase, and 7.20 µL T4 PNK. (15) Repaired RNA fragments were reverse transcribed by adding 100 nL of RT mix per well and incubation at 50 °C for 1 h. Per plate, the RT mix contains 38.7 µL Nuclease-free water, 3 µL 0.1 M DTT, 4.5 µL 1 M KCl, 3 µL 50 % PEG8000, 0.3 µL 20 mg/mL BSA, 4.5 µL 10 mM dNTPs, and 6 µL SuperScript III. (16) DNA fragments were blunt-ended by adding 150 nL DNA repair mix per well and incubation for 30 min at 37 °C, followed by 20 min at 75 °C for enzyme inactivation. Per plate, the DNA repair mix contains 54.6 µL Nuclease-free water, 1.2 µL 5 M NaCl, 10.5 µL 1 M Tris with pH 7.5, 1.5 µL 1 M MgCl_2_, 4.5 µL 0.1 M DTT, 4.5 µL 50 % PEG8000, 0.45 µL 20 mg/mL BSA, 3.9 µL 100 mM ATP, 1.95 µL 10 mM dNTPs, 3 µL TL Exo1, 1.95 µL Klenow large fragment, and 1.95 µL T4 PNK. (17) Blunt fragments were subsequently A-tailed by adding 150 nL A-tailing mix per well and incubation for 15 min at 72 °C. Per plate, the A-tailing mix contains 76.7 µL Nuclease-free water, 6 µL 1 M KCl, 4.5 µL 50 % PEG8000, 0.45 µL 20 mg/mL BSA, 1.2 µL 100 mM dATP, and 1.2 µL Taq360. Next, fragments were ligated to forked adapters containing T-overhangs (see supplementary table 2 for sequences). (18) For adapter ligation, 50 nL of 5 mM adapters in 50 mM Tris with pH 7 was added to each well with the Mosquito HTS (ttp labtech). (19) After centrifugation, 150 nL of adapter ligation mix was added and (20) plates were incubated for 20 min at 4 °C, followed by 16 h at 16 °C for ligation and 10 min at 65 °C for ligase inactivation. Per plate, the adapter ligation mix contains 17.1 µL Nuclease-free water, 2.1 µL 1 M MgCl_2_, 2.7 µL 1 M Tris with pH 7.5, 40.5 µL 0.1 M DTT, 4.5 µL 50 % PEG8000, 0.45 µL 20 mg/mL BSA, 2.1 µL 100 mM ATP, and 21 µL 400.000 U/mL T4 DNA Ligase. (21) Before pooling, 1 mL of Nuclease-free water was added to each well to minimise material loss. (22) Ligation products were pooled by centrifugation into oil-coated VBLOK200 Reservoirs (ClickBio) at 500 g for 2 min and (23) the liquid phase was transferred into 1.5 mL Eppendorf tubes and (24) was purified by centrifugation at 13000 g for 1 min and transferred into a fresh tube. Steps 23 and 24 were performed twice. (25) DNA fragments were purified using Ampure XP beads (Beckman Coulter - prediluted 1 in 8 in bead binding buffer – 1 M NaCl, 20 % PEG8000, 20 mM Tris with pH 8, 1 mM EDTA) at a bead-to-sample ratio of 0.8. (26) After 15 min incubation at RT, beads were washed twice with 1 mL 80 % ethanol, resuspending the beads during the first wash and (27) resuspension in 17 µL Nuclease-free water. After 2 min elution, the resuspended beads were transferred into a fresh 0.5 mL tube. (28) Second-strand synthesis was performed per plate using the NEBNext^®^ Ultra™ II Non-Directional RNA Second Strand Synthesis Module. For this, 2 µL 10x reaction buffer was mixed with the sample on ice, followed by addition of 1 µL enzyme, gentle mixing, and incubation at 16 °C for 2.5 h and 65 °C for 20 min. (29) A second cleanup was performed after second strand synthesis by adding 26 µL of undiluted Ampure XP beads. The beads were resuspended in 8 µL Nuclease-free water. (30) After the cleanup, the DNA was linearly amplified by in vitro transcription with 12 µL of the MEGAscript™ T7 Transcription Kit (Fisher Scientific, AMB13345) for 12 h at 37 °C. (31) The produced RNA was purified using RNA Clean XP beads (Beckman Coulter) with a 1.3x beads-to sample ratio (26 µL beads), followed by sample resuspension in 12 µL Nuclease-free water. (32) At this stage, 6 µL aRNA can be stored at −80 °C, while continuing with the remaining 6 µL. Next, rRNA was depleted using 2 µL Hybridisation Buffer (1000 mM NaCl in 500 mM Tris with pH 7.5) and 2 µL 25 µM rRNA depletion oligos (supplementary table 3). (33) Oligos were hybridised to their rRNA targets by incubation at 95 °C for 2 min, followed by a gradual temperature decrease of 0.1 °C/s down to 45 °C. Once at 45 °C, 10 µL RNAse mix (2 µL thermostable RNAseH (Epicentre), 8 µL RNAse buffer: 250 mM NaCl and 50 mM MgCl_2_ in 125 mM Tris with pH 7.5) at 45 °C was added and samples were incubated at 45 °C for another 30 min. (34) DNA template and unused depletion probes were removed through addition of 4 µL RNase-free DNAse (Promega) with 5 µL 10 mM CaCl_2_ in 30 µL Nuclease-free water and incubated at 37 °C for 30 min. (35) RNA was purified using 80 µL RNA Clean XP beads per sample and elution in 6 µL water. (36) 5 µL supernatant was transferred to a new 0.5 mL Eppendorf tube for RNA adapter ligation. For this, 1 µL 20 µM RA3 (/5rApp/TGGAATTCTCGGGTGCCAAGG/3SpC3/) was added, mixed and incubated for 2 min at 70 °C, followed by cooling on ice. On ice, 4 µL RNA ligation mix was added, mixed by pipetting and incubation for 1 h at 25 °C (Lid temperature should not exceed 25 °C). Per sample, RNA ligation mix contains 1 µL T4 RNA ligase 2, truncated (NEB), 1 µL T4 RNA ligase reaction buffer (NEB), 1 µL RNAseOUT (Invitrogen), and 1 µL Nuclease-free water. (37) RNA was primed for reverse transcription by adding 1 µL 10 mM dNTPs and 2 µL 20 mM reverse transcription primer - RTP (GCCTTGGCACCCGAGAATTCCA) and (38) hybridization through incubation at 65 °C for 5 min and direct cool-down on ice. (39) Reverse transcription was performed by addition of 4 µL first strand buffer (Invitrogen), 1 µL 0.1 M DTT, 1 µL RNaseOUT (Invitrogen), 1 µL SuperscriptIII (Invitrogen) and 1 µL Nuclease-free water and (40) incubating the mixture at 50 °C for 1 h followed by 15 min at 70 °C. (41) Single-stranded DNA was purified using 1 µL RNAseA (Thermo Fisher) and incubation for 30 min at 37 °C. (42) DNA was cleaned up with 22 µL Ampure XP beads and resuspension in 20 µL Nuclease-free water. 10 µL can be stored at −80 °C. (43) The library PCR to add the Illumina small RNA barcodes and handles was performed by adding 25 µL NEBNext Ultra II Q5 Master Mix (NEB, M0492L), 11 µL Nuclease-free water and 2 µL RP1 and RPIx primers each (10 mM, for details see Illumina small RNA indexing primer sequences) to the 10 µL single-stranded DNA product. (44) PCR incubation: activation for 30 sec at 98 °C, 8 cycles 10 sec at 98 °C, 30 sec at 60 °C, 30 sec at 72 °C, final amplification 10 min at 72 °C (45) PCR products were cleaned by 2 consecutive DNA bead clean-ups with a 0.8x bead-to-sample ratio, eluting in 30 µL Nuclease free water after the first round and (46) in 11 µL for the final product. Abundance was determined using a dsDNA hs QUBIT, and after adjusting concentrations to ∼2 ng/µL quality was assessed using a DNA hs bioanalyzer.

### pA-MN production

The pA-MN fusion protein was produced following the methods section in ^66^.

### Sequencing

Final DNA libraries are sequenced paired-end 100 bp, on either a Novaseq or NextSeq2000.

### Transcriptome analysis

Fastq files were processed into .loom files using the T-ChIC Snakemake ^67^ (see https://github.com/marloes3105/tchic/tree/main/workflows/1.snakemake-workflow/). The pipeline works as follows: reads were demultiplexed for CEL-seq2 barcodes using SingleCellMultiOmics’ (SCMO) demux.py (v0.1.30), polyA tails were trimmed with SCMO trim_vasa.py and adapters were trimmed with cutadapt ^68^ (v4.1), followed by mapping of the reads with STAR ^69^ (v2.5.3a) to the 129/B6 SNP-masked GRCm38 mouse genome (Ensembl 97) for the gastruloid data and GRCh38 (Ensembl 97) for the K562 cell line data. Reads were tagged and deduplicated using SCMO bamtagmultiome.py, and velocyto ^70^ (v0.17.17) was run to generate output .loom files. Downstream analysis was performed with Scanpy ^71^ (v1.8.1). Cells with less than 1.000 or more than 50.000 reads, and less than 100 genes detected were filtered out. Cells with more than 0.6 percent mitochondrial reads were also excluded. Mitochondrial reads, Malat1, SnoRNAs, and ribosomal reads were excluded from further analysis. Counts were normalised to 1.000 transcripts per cell and logarithmized. The number of counts, percentage of mitochondrial reads, and ChIC HPTM profiled for the same cell were regressed out and each gene was scaled to unit variance with a maximum of 10.

Principal component analysis was performed and used to filter out cells with low ChIC quality (see cell filtering). Neighbours were calculated with sc.pp.neighbors with parameters n_neighbors = 5, n_pcs = 50, metric = manhattan and the UMAP was generated with sc.tl.umap with parameters min_dist = 0.4, spread = 0.6. Clustering was performed using sc.tl.leiden, with the resolution set at 0.04 for germ layer separation. For detailed cell typing, germ layers were separated and re-clustered with resolutions set at 0.3 for haemogenic, 0.2 for endoderm, 0.4 for ectoderm, 0.8 for mesoderm, and 0.9 for pluripotent, followed by labelling and merging of the full dataset. Clear separation of marker genes was leading for the determination of cluster sizes and boundaries.

### ChIC analysis

Fastq files were processed into count tables using the T-ChIC Snakemake ^67^ (see https://github.com/marloes3105/tchic/tree/main/workflows/1.snakemake-workflow/). The pipeline works as follows: reads were demultiplexed for T-ChIC barcodes with a hamming distance of 0 using SingleCellMultiOmics’ (SCMO) demux.py (v0.1.30), followed by adapter trimming using cutadapt ^68^ (v4.1). Reads were mapped paired-end with bwa mem ^72^ (v0.7.16a) to the 129/B6 SNP-masked GRCm38 mouse genome (Ensembl 97) with the following parameters: -M -I 1000. Reads were tagged with SCMO bamtagmultiome.py using a blacklist for specific genomic regions ^73^ and duplicates and reads with an insert size above 1000, mapping quality below 50, or non-TA ligation motif were removed.

To exclude low quality cells, transcript-based PCs were used to determine neighbourhoods of comparable cells. The local chromatin enrichment of each cell was compared to the neighbourhood average. To determine the neighbourhood average, pseudo-bulks of total count normalised cells were created and the highest covered bins containing 80% of the reads were determined. Next, the read fraction of the cell of interest in those neighbourhood-enriched bins was determined and cells with an enrichment of 2 std below average were marked as low quality. By definition, the enrichment distribution is around 80%. The script that was used is cellSelection_PCA.py with parameters: -neighbors *number of neighbors to use* -cutoff *minimal number of reads* -chic_csv *genome wide ChIC count table/merged_count_table_50000.csv* -trans_csv *PCs exported from scanpy* -o *folder name for output*. For all datasets, 200 neighbours were used, and minimum reads per cell were set at 300 for H3K4me3 and 900 for H3K27me3.

### Clustering and visualization of ChIC data

For clustering of cells based on scChIC signal, the UMI counts were aggregated in 50kb genomic bins using function scCountReads from sincei package ^74^ (v0.4), with parameters ‘-bs 50000 --minMappingQuality 10 --samFlagInclude 64 -bl blacklist.bed’ followed by latent schematic analysis (LSA) of cells*bin matrix yielding Cell*Topic and Region*Topic matrices, as described before ^75^. Next, we used scanpy to calculate 30 nearest neighbors using the 50 topics from the LSA output (dropping Topic 1, which strongly correlates with read depth), and used it to build the paga graph using scanpy.tl.paga with ‘threshold = 0.05’. UMAPs were initialized using the PAGA graph using ‘scanpy.tl.umap(’min_dist=0.1, spread=1’, init_pos=’paga’)’. Multi-UMAP visualization of cells based on their ChIC and RNA information was performed using the plotMultiUMAP function in the sctoolbox R package (https://github.com/vivekbhr/sctoolbox, v0.1). To plot the ChIC and RNA signal over genomic tracks, we ran scCountReads with the same parameters as above for the selected genomic regions, but at a binsize of 5 kb (-bs 5000), and re-exported the resulting count table into the “bedgraphmatrix” format. Dual (ChIC+RNA) signal was exported after superimposing the ChIC counts over the RNA counts. The exported counts, along with the bigwigs (representing the sum of counts) were plotted using pyGenomeTracks ^76^ (v 3.7).

### Comparison with other protocols

The chromatin and transcriptome count tables for published protocols were downloaded from their GEO repositories ^29,30,32^. To make the data biologically comparable, we only selected ES cells ^29,30,32^. For the chromatin fraction, Rang *et al.* data contained counts aggregated in 10 kb genome-wide bins. We therefore aggregated the chromatin counts from all other studies (including ours) on the same bins. For the transcriptome, we used the gene-level UMI counts from all studies. To compare the variance in chromatin counts based on ranked gene expression, we ordered genes by their mean expression across all cells in their respective study and divided them into 10 fractions. We then overlapped the gene promoters (TSS +-1kb) with the chromatin bins, summed up the counts in overlapping bins per gene, and plotted the average, log-scale counts per gene across cells in the respective study.

### Pseudotime determination

Pseudotime was calculated using cellrank’s ^77^ (v1.1.0) pseudotime embedding in combination with scvelo ^78^ (v0.2.4). In short, terminal states were calculated based on lineage + day clusters (‘lin_day’) using cr.tl.terminal_states with parameter weight_connectivities = 0.2, and initial states were calculated with cr.tl.initial_states. Lineages were computed with cr.tl.lineages. Finally, pseudotime was calculated with scv.tl.recover_latent_time with parameters root_key = ‘initial_states_probs’ and end_key = ‘terminal_states_probs’.

### Classification of gene states per cell type

To determine cell type-specific gene states, pseudobulks were generated by randomly subsampling 400 cells processed for each HPTM (H3K4me3 and H3K27me3). To classify the resulting count data into binary states (negative/positive), we implemented an unsupervised Gaussian Mixture Model (GMM) approach using two components. This method allowed us to define a data-driven threshold for binarization without imposing a fixed cutoff. A two-component GMM was fitted to the data using the scikit-learn implementation ^79^ (GaussianMixture, v1.3.2). The model assumes the observed distribution arises from a mixture of two Gaussian distributions, representing “background” and “positive” states.

To identify and remove statistically insignificant changes between cell types, we employed a one-way Analysis of Variance (ANOVA) combined with fold-change filtering, using 5 replicas generated by subsampling to estimate within-cell type variance. A one-way ANOVA test was performed independently for each gene using scipy.stats.f_oneway ^80^ (v1.10.1) using the formula:

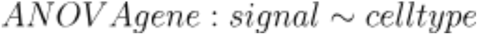

Genes with -log10 p-value smaller than 2.3 for RNA and H3K4me3 and smaller than 5 for H3K27me3, as well as log_2_ fold change smaller than 2 for RNA and H3K4me3 and smaller than 3 for H3K27me3 were considered as stable. For those genes, the relevant binarisation result was replaced with the dataset median.

### Classification of gene states along differentiation trajectories

To determine gene states along differentiation trajectories, pseudobulks were generated by separating cells based on their lineage. Epiblast and PGC-like cells were excluded for this analysis. Cells were sorted along pseudotime and pseudobulks were generated of 50 non-overlapping cells for both HPTMs and of 100 non-overlapping cells for RNA.

To classify the resulting count data into binary states (negative/positive), we implemented an unsupervised Gaussian Mixture Model (GMM) approach using two components. To avoid noise-related frequent transitions between states, the count data was smoothed using a rolling mean with a stepsize of 10 for RNA and 15 for both HPTMs. A two-component GMM was fitted similar to the cell type data. For the binarisation, each pseudobulk was only used once, to avoid over-representation of shared early time points.

To identify and remove statistically insignificant changes, we performed two ANOVA tests: one for pluripotent cell types including caudal epiblast, and one for more mature cell types. We used the more significant result from both tests. For all modalities, genes with - log10 p-value smaller than 2.3 as well as log_2_ fold change smaller than 2 were considered as stable. For those genes, the relevant binarisation result was replaced with the dataset median. For state transitions the median of pseudobulk steps 1-10 was used as mESC state, steps 50-60 (representing the caudal epiblast) as pluripotent and the last 10 steps per trajectory as differentiated cell types.

### Identification of differential H3K4me3-marked and transcribed TSS

For analysing differential usage of annotated TSS, for each gene all TSS that are at least 10 kb downstream of a previous TSS of the same gene, as well as additional TSS less than 10 kb from the end of the gene, were selected from Mus_musculus.GRCm38.97.gtf (ensembl.org) to avoid overlap. Single-cell signals were counted in a 2.5 kb upstream to 7.5 kb downstream window of the TSS for H3K4me3 and 10 kb downstream of the TSS for RNA. Cells were subsampled to generate 5 equally sized non-overlapping pseudobulks per lineage. To classify the resulting count data into binary states (negative/positive), we implemented an unsupervised GMM approach with two components to identify H3K4me3-positive TSS, transcribed TSS and transcribed genes, using per-TSS H3K4me3 counts, per-TSS RNA counts, and whole-gene transcript counts, respectively. The median of the state obtained for the 5 replicas was used. To identify lineage-specific H3K4me3 enrichments at individual TSS, we performed a generalized linear model (GLM) analysis with a Poisson regression framework using statsmodels ^81^ (v0.14.1). For each TSS, H3K4me3 counts were modeled as a function of lineage identity using the following formula:

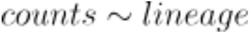

To ensure numerical stability and avoid singular model fits, small Gaussian noise (μ = 0, σ = 0.1) was added to the counts when all values were identical within a region and TSS with zero variance across all replicates were assigned neutral statistics (p=1). Finally p-values from the GLMs were corrected for multiple hypothesis testing using the Benjamini–Hochberg false discovery rate (FDR) procedure. To identify differential TSS transcription across lineages, we used a similar GLM framework incorporating interaction terms between lineage and TSS-level expression. This allowed us to evaluate whether lineage-specific changes in TSS signal diverged significantly from whole-gene expression levels with the following formula:

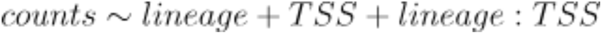

Where counts are log-transformed RNA counts (with a pseudocount of 1 and jitter of 0-0.1 to improve numerical stability). TSS were considered differentially expressed or differentially H3K4-methylated with a -log10 p-value of > 2.3 and a log_2_ fold change of >1.

### Identify correct TSS to use per gene

For genes with multiple annotated TSS that do not show H3K4me3 or RNA signal in any lineage, the primary annotated TSS was used for further analysis. For genes with only H3K4me3 signal but no transcription, the TSS with the largest H3K4me3 signal over all lineages was used. For genes with recovery of both H3K4me3 and RNA signals, the first TSS along the gene that was H3K4me3- and RNA-positive was used. For RNA, the threshold was set at 10% of the maximal TSS-specific RNA counts of the gene.

### Lorentz plots

Lorentz plots were based on genome-wide 50 kb count tables from lineage-specific pseudobulks of 1400 cells each. Bins were sorted per lineage from lowest to highest counts and the cumulative sum was plotted on the y-axis relative to the number of genomic bins summed over.

### Identifying germ layer-specific differentially expressed genes

To identify genes that become expressed specifically in one of the three germ layers, differential gene analysis was performed using DEseq2 ^52^ on the transcriptome dataset. First, cells were divided into four groups: pluripotent, ectoderm, mesoderm and endoderm (see Figure 6A). To focus specifically on genes that change in a lineage specific-manner, we limited the analysis on cells after the Wnt activation pulse, excluding Epiblast, PGC-like cells and mESCs. The pluripotent cluster was compared sequentially with the three differentiated clusters. Decoupler ^82^ (v1.9.2) was used to determine and filter pseudobulks, using plate numbers as replicates. Briefly, cells were aggregated into pseudobulks using dc.get_pseudobulk, with groups_col = [‘pluripotent’, ‘differentiated’], sample_col = plate number, and mode = ‘mean’. Genes with < 250 total counts or expressed in < 5 pseudobulks were removed using dc.plot_filter_by_expr. Pseudobulks with < 300.000 total raw counts or < 50 cells for mesoderm and ectoderm, and < 20 cells for endoderm were removed. Counts were processed using scanpy ^71^ (v1.10.3). Pseudobulk counts were normalised using sc.pp.normalize_total with target_sum = 1e4, log-transformed with sc.pp.log1p, and scaled using sc.pp.scale with max_value = 10. The adata with stored pseudobulks was used to build the deseq2 object with DeseqDataSet from pydeseq2 ^83,84^ (v0.5.0). Log fold changes were calculated using dd.deseq2(), and log_2_ fold changes and adjusted -log10 p-values were extracted. Genes with a -log10 p-value of < 0.05 and log_2_ fold change of > 2 were identified as differentially expressed, and overlapping genes between the three sets were removed.

### Determining the point of significant change along pseudotime

Cells were grouped into pseudobulks along the 7 trajectories and for each pseudobulk, the rolling mean average per gene and modality was used (see “Classification of gene states along differentiation trajectories”). For each gene and modality, the standard deviation of these values was calculated across all trajectories using numpy’s np.std ^85^ (v2.0.2). The lower and upper thresholds for significant change were calculated by adding or subtracting, respectively, 2x standard deviation from the value of the first pseudobulk. For each gene, modality, and trajectory, the first pseudobulk to cross the lower or upper threshold got assigned a state of - 1 or +1, respectively. For RNA and H3K4me3, only pseudobulks changing to +1 were stored (significant increase). For H3K27me3, only pseudobulks changing to −1 were stored (significant decrease). For scatterplots and violin plots, each gene in each trajectory was required to change in both RNA and the displayed HTPM. For violin plots, the step number of the pseudobulk with significant change was used, and the RNA step was subtracted from the HPTM step.

### Code and data availability

All code used and annotated notebooks for performing the analysis and generating figures can be found on github (https://github.com/marloes3105/tchic, release v1.0).

## Acknowledgements

We thank Samy Kefalopoulou and Vladyslav Bondarenko for critical reading of the manuscript. pK19pA-MN was a gift from Ulrich Laemmli (Addgene plasmid 86973, http://n2t.net/addgene:86973; RRID:Addgene_86973). The 129S1/SvImJ / C57BL/6 embryonic stem cell line was a gift from Matyas Flemr and Marc Bühler. We thank Reinier van den Linden and the Hubrecht Flow Cytometry Core facility for their essential help with experiments. We acknowledge the Utrecht Sequencing Facility (USEQ) for providing sequencing service and data. USEQ is subsidized by the University Medical Center Utrecht and The Netherlands X-omics Initiative (NWO project 184.034.019). This work was supported by European Research Council Advanced grant ERC-AdG 101053581-scTranslatomics. The HFSP (LT000209/2018-L), Marie Skłodowska-Curie Actions (798573) and ERC starting grant 101164799-scEpiTarget supported P.Z. The EMBO LTF (ALTF 1197–2019) supported V.B. This work is part of the Oncode Institute, which is partly financed by the Dutch Cancer Society. The funders had no role in study design, data collection and analysis, decision to publish or preparation of the manuscript.

## Author contributions

M.B., P.Z. and A.v.O. designed the project; M.B., P.Z. and F.S. developed experimental methods; M.B. and P.Z. performed experiments; M.B. and B.A.d.B. developed the T-ChIC demultiplexing and preprocessing pipeline. M.B., P.Z. and V.B. analysed the data. M.B. and P.Z., wrote the manuscript with input from B.A.d.B., V.B., and A.v.O.

## Supplemental figures

**Figure S1.**
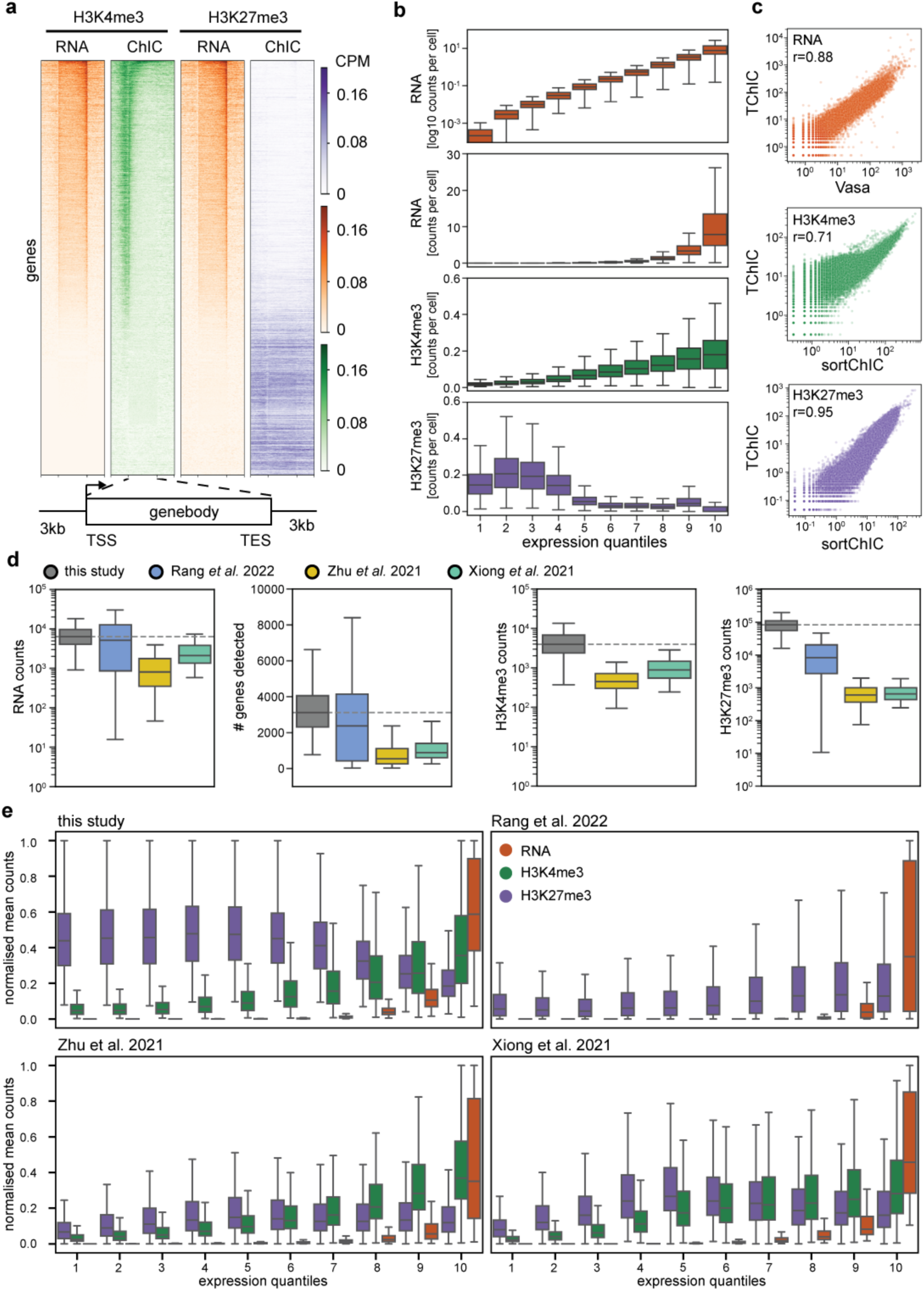
T-ChIC recovers biologically expected chromatin distributions and outcompetes alternative methods in sensitivity and selectivity. a) Heatmap of read distribution in a window of 3 kb around protein-coding gene bodies. Pseudo-bulk results from one plate of K562 cells per HPTM are shown. b) Boxplots of single-cell read averages over genes split into quantiles of log10 expression. From top to bottom, the following measurements are shown: RNA log10 counts (orange), RNA counts (orange), H3K4me3 (green) and H3K27me3 (purple). c) Scatter plots showing Pearson correlation of T-ChIC to VASA-seq and published ChIC results for K562 cells. shown are RNA (top, orange), H3K4me3 (middle, green) and H3K27me3 (bottom, purple). d-e) Comparison of RNA and chromatin sensitivity (d) and selectivity (e) with published single-cell methods that allow simultaneous measurement of transcription and HPTM distribution. Rang et al. is based on DamID, which uses transgenic cell lines expressing Dam fusion proteins in combination with life cell sorting into 384 well plates. Zhu et al. and Xiong et al. are based on cut&tag (pA-Tn5) and the library prep is performed by combinatorial barcoding. d) From left to right, boxplots of uniquely mapped transcript reads, number of genes with at least one read, uniquely mapped H3K4me3 and H3K27me3 reads are shown. e) Boxplots of single-cell read averages over genes split into quantiles of log10 expression are shown for RNA (orange), H3K4me3 (green) and H3K27me3 (purple). Values are scaled per modality and one plot is shown per study.

**Figure S2.**
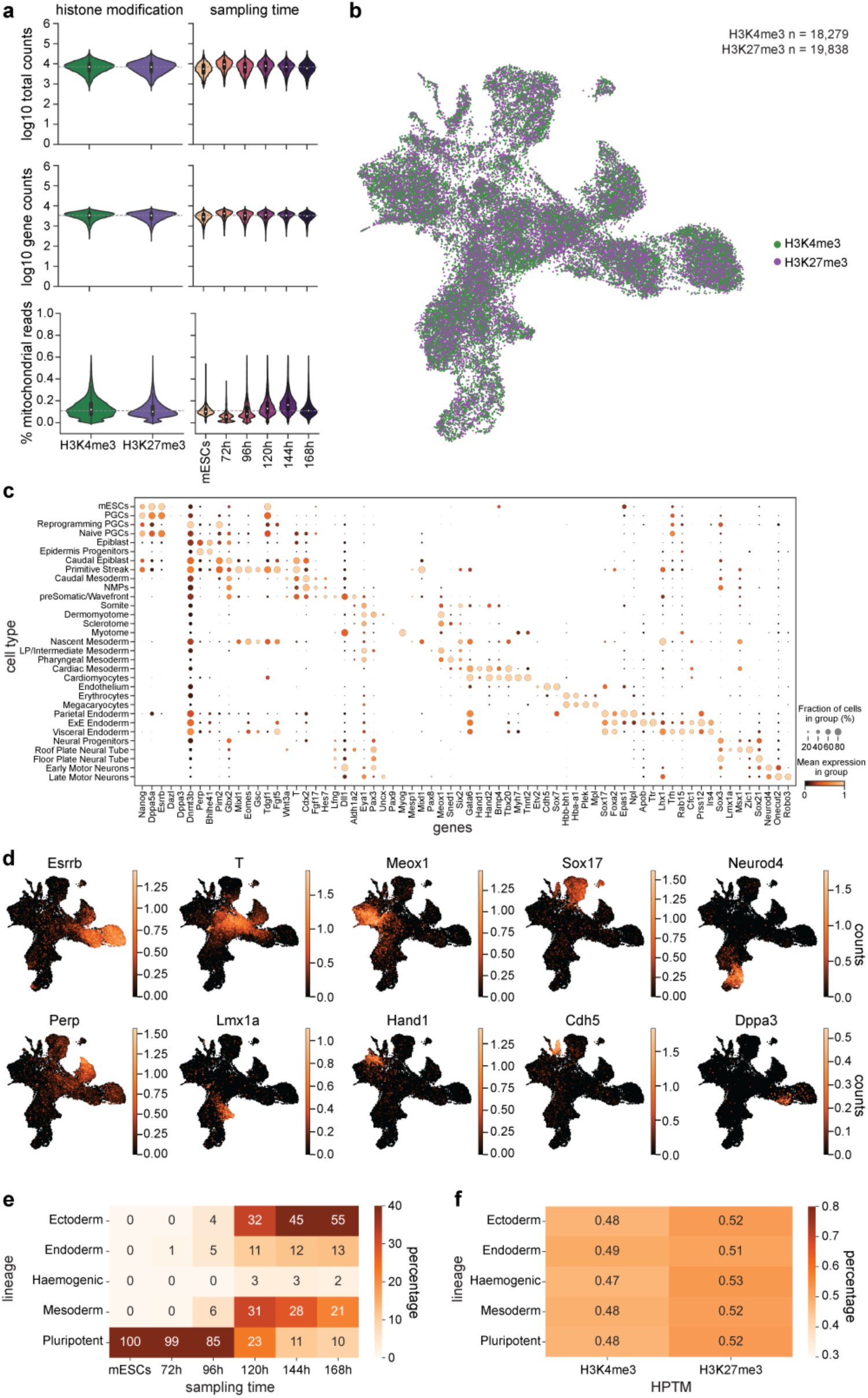
Gastruloid time course recovers differentiation time point-dependent cell types. a) Transcriptome QC split by HPTM profiled simultaneously (left) or sampling time point (right). The following QC metrics are shown: log10 values of total counts per cell (top), log10 values of total genes recovered per cell (middle), and percentage of mitochondrial reads per cell (bottom). b) Transcriptome UMAP coloured by the HPTMs profiled together with RNA: H3K4me3 (green, n = 18,279) and H3K27me3 (purple, n = 19,838). c) Dot plot showing expression of marker genes used for cell type annotation in Figure 2C. Dot colour represents the mean expression of marker genes per specified cluster, and dot size represents the fraction of cells within the group that show expression of the gene. d) Transcriptome UMAPs showing expression levels and patterns of selected marker genes: *Esrrb*, pluripotency; *T*, symmetry breaking and elongation factor; *Meox1*, early mesoderm; Sox17, endoderm; *Neurod4*, ectoderm; *Perp*, epiblast; *Lmx1a*, neural tube; *Hand1*, heart progenitors; *Cdh5*, endothelium; *Dppa3*, PGC-like. e) Percent of lineages recovered for each sampling time point. f) Percent of lineages recovered for each chromatin modification profiled.

**Figure S3.**
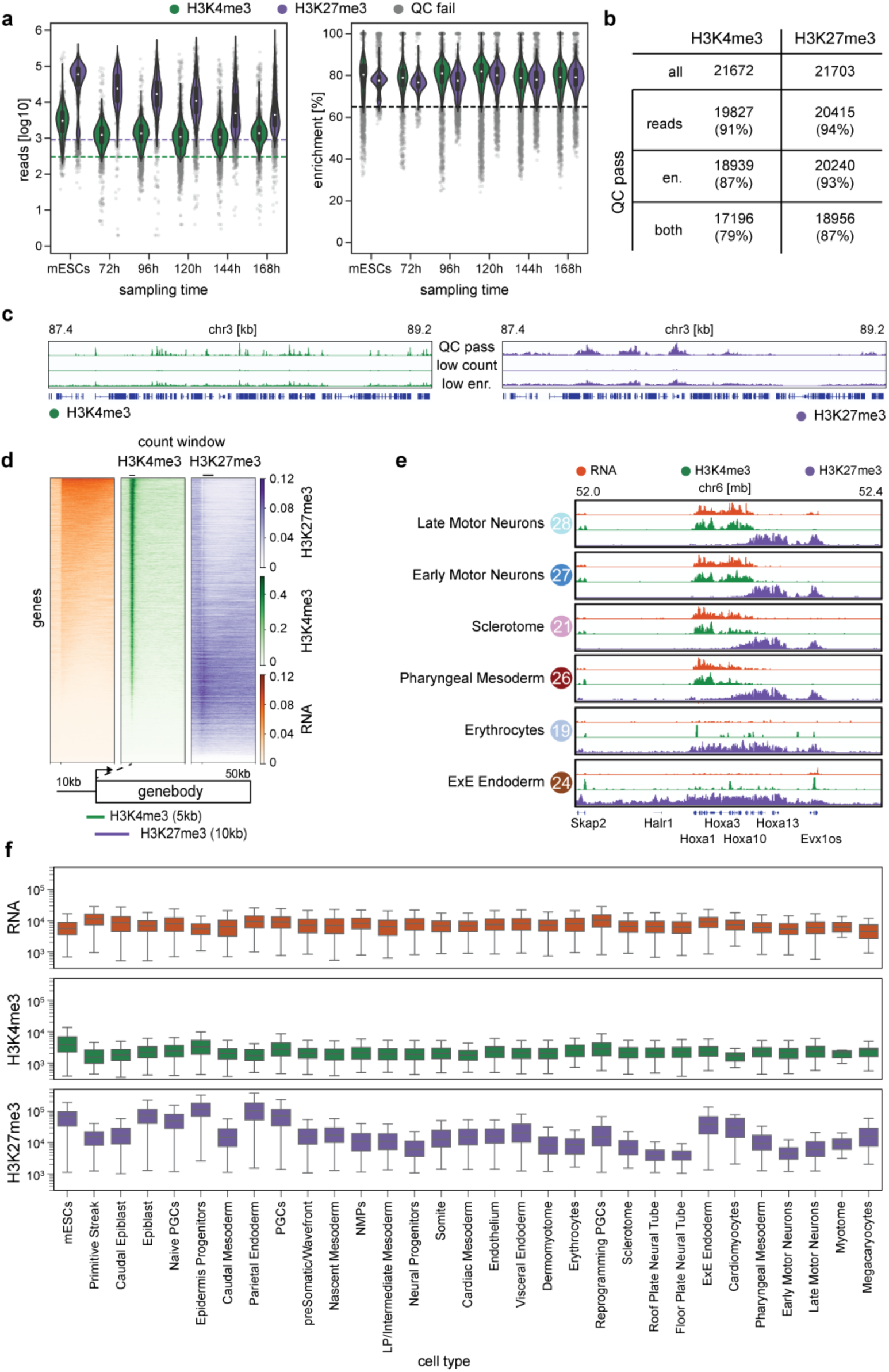
QC filtered T-ChIC data recovers cell type-specific chromatin changes in gastruloids. a) Chromatin QC split by sampling time point and HPTM profiled. Left: log10 uniquely mapped reads per cell, right: fraction of reads assigned to genomic bins enriched in local neighbourhoods. b) Number and percentage of cells passing QC filtering. c) Coverage plots showing the signal obtained from 1000 randomly selected cells passing QC, or failing due to low reads or enrichment. d) Heatmap of read distribution 10 kb upstream and 50 kb into the gene body of gastruloid pseudo-bulk data. RNA (orange, left), H3K4me3 (green, middle) and H3K27me3 (purple, right) are shown. Lines on top indicate counting windows for gene-based analysis. e) Pseudo-bulk representation per cluster of single-cell coverage plots shown in Figure 3C. A 400 kb window on chromosome 6 covering the Hoxa cluster is shown. f) Log10 uniquely mapped reads per cell are shown, split per cell type for RNA (orange, top), H3K4me3 (green, middle) and H3K27me3 (purple, bottom).

**Figure S4.**
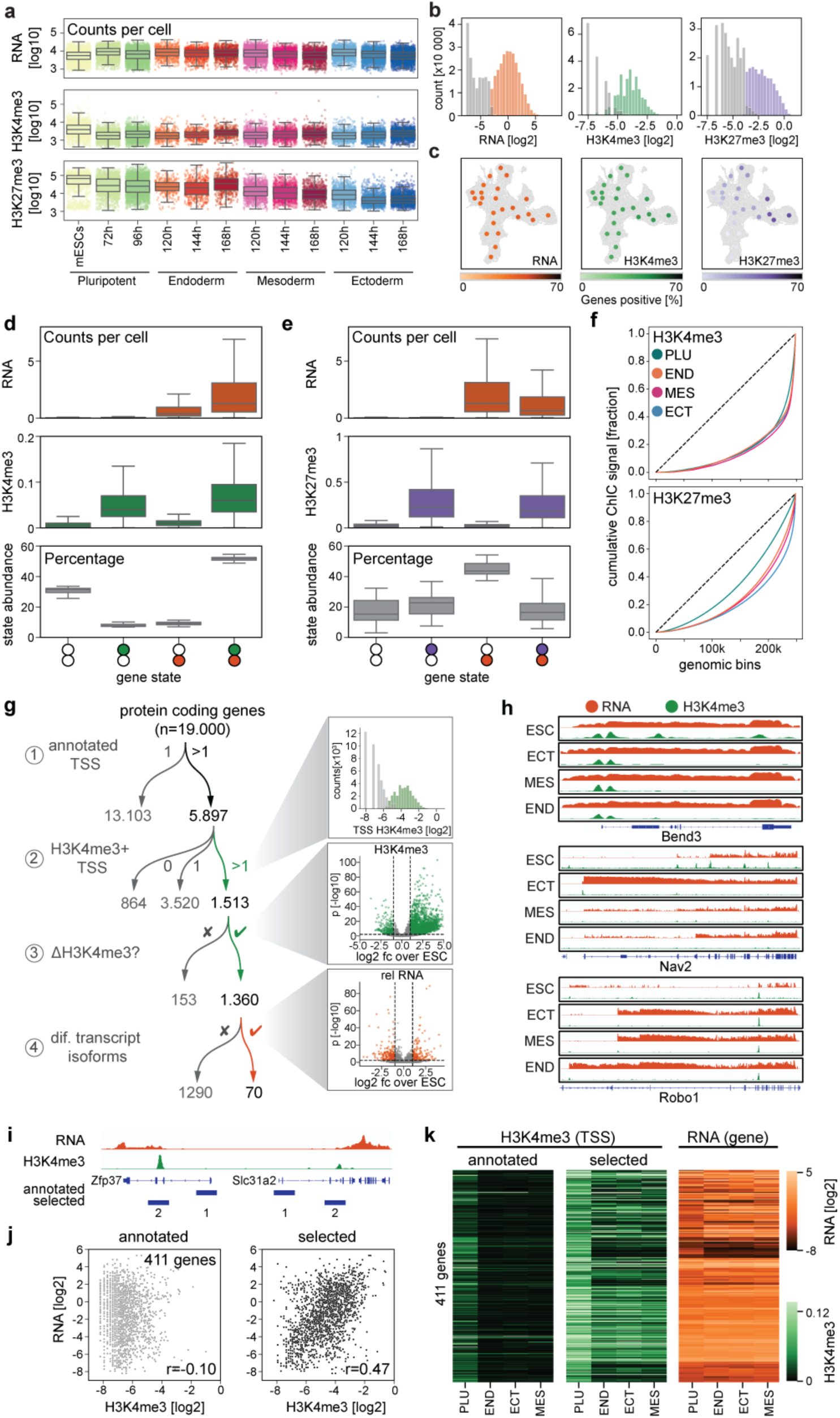
The exit of pluripotency is characterised by a global decrease in H3K27me3 and H3K4me3 and a switch in transcription start site (TSS) usage. a) Boxplot of total counts per modality and in cells belonging to indicated lineage and harvesting timepoints. b) Histograms of gene-wise log_2_ RNA (left), promoter-based H3K4me3 abundance (middle, 2.5- and 7.5kb+ of the TSS) and H3K27me3 abundance (right, 2.5- and 7.5kb+ of the TSS). Genes are separated based on a mixture model into negative (grey) and transcribed (orange), H3K4me3 methylated (green), or H3K27 methylated (purple). c) UMAPs with pseudobulk dots coloured by the fraction of genes classified as positive for the indicated layer. d-e) Boxplots showing distribution of RNA counts and H3K4me3 (d) or RNA and H3K27me3 counts (e) as well as fraction of genes falling into each of the four gene states. RNA and chromatin counts are plotted per gene and cell type. Abundance is plotted per cell type. f) Lorentz plot, showing the genome-wide distribution of H3K4me3 (top) and H3K27me3 (bottom) in mESCs and the three germ layers. g) Graphical summary of the different analysis steps used to distinguish genes with multiple annotated TSS (1), in case of multiple annotated TSS have at least two covered by H3K4me3 (2), in the case of multiple marked TSS per gene, which TSS are differentially marked between lineages (3) and for differentially marked genes, if TSS used for transcription start changes between lineages (4). This allows the identification of marked, differentially marked and differentially used TSS. h) Pseudobulk coverage plots displaying RNA (orange) and H3K4me3 (green) coverage of three loci spanning example genes Bend3 (top), Nav2 (middle) and Robo1 (bottom) displaying differential H3K4me3 TSS coverage without (Bend3), or with (Nav2, Robo1) corresponding TSS usage switch from pluripotent cells to the three lineages. i) Coverage tracks of RNA and H3K4me3 of two genes where the alternative annotated TSS is used. Bottom tracks indicated the annotated major TSS and the selected used TSS. j) Scatterplot showing the gene-wise relationship between H3K4me3 and transcription for the main annotated TSS left and the selected used TSS right for the 411 genes where the annotated TSS is not used in the gastruloid system. k) heatmap showing H3K4me3 over annotated and selected TSS on the left and the corresponding gene level expression on the right.

**Figure S5.**
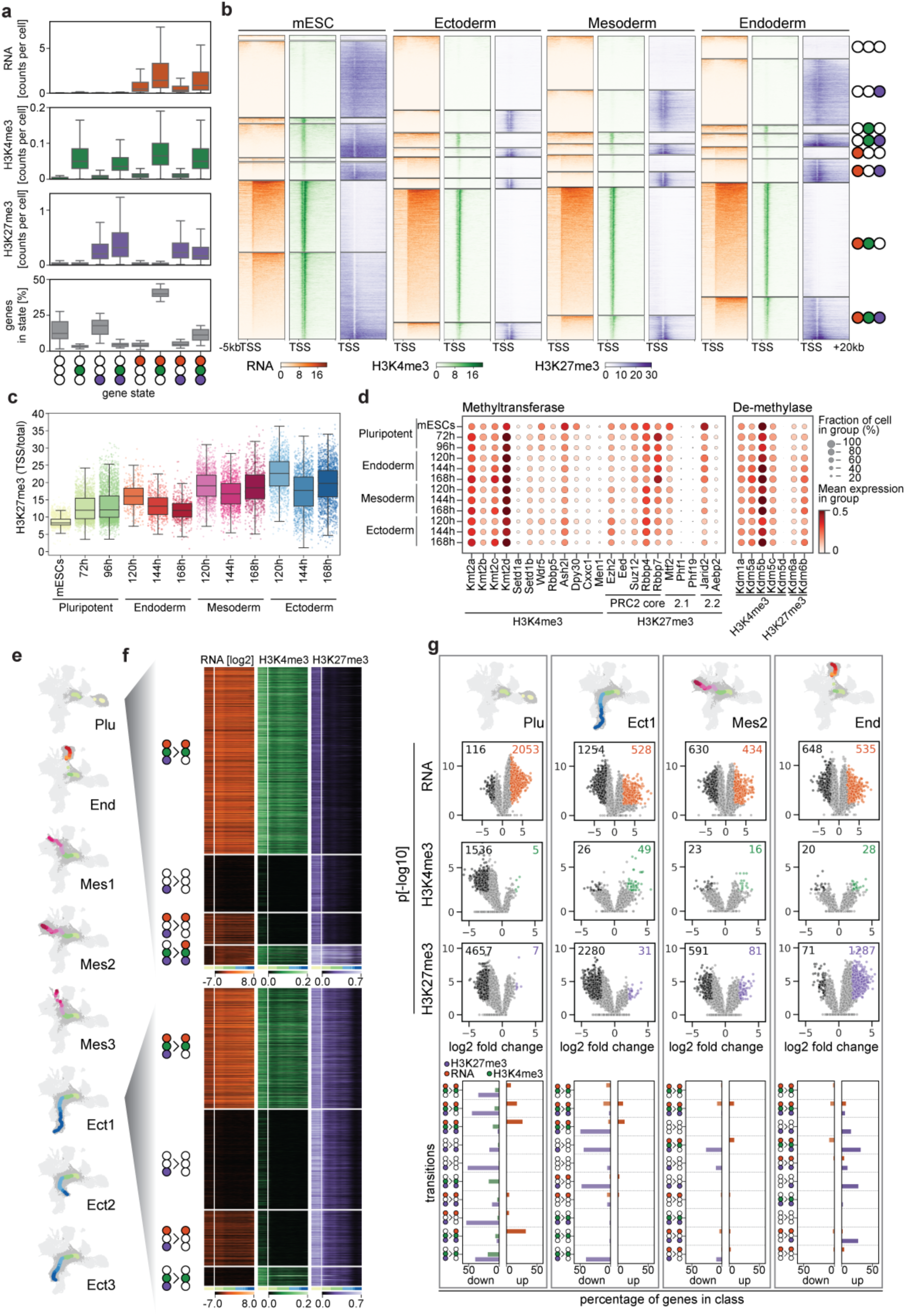
Classification of genes into eight activity states recovers germ layer-specific loss of H3K27me3. a) Box plots showing distribution of RNA counts, H3K4me3 and H3K27me3 counts as well as fraction of genes falling into each of the eight gene states. RNA and chromatin counts are plotted per gene and cell type. Abundance is plotted per cell type. b) Cover plots showing RNA (orange), H3K4me3 (green) and H3K27me3 (purple) distribution over the TSS of individual genes separated into the eight gene states. c) Boxplot showing the fraction of total H3K27me3 fragments per cell falling into promoter regions (2.5 kb upstream to 7.5 kb downstream of the TSS) split by sampling time and germ layer. d) Dotplot showing expression of key epigenetic regulators of the indicated epigenetic modifications. e) UMAPs indicating the 8 isolated trajectories. Small dark grey dots represent single cells included in the trajectory, large dots coloured by lineage and sampling time represent generated pseudobulks. f) Heatmaps showing RNA expression (orange), H3K4me3 (green) and H3K27me3 (purple) coverage of gene sets separated by state changes across pseudotime for indicated trajectories (top: pluripotency, bottom: ectoderm-1). g) Top: UMAPs of four example trajectories for pluripotency and the three lineages. Middle: volcano plots indicating significant up- or down-regulation of genes in indicated trajectories across the RNA (top: black/orange), H3K4me3 (middle: black/green) and H3K27me3 (bottom: black/purple) modalities. Bottom: bar plots summarising significant up- or down-regulation of genes across the three modalities (orange, RNA; green, H3K4me3; purple, H3K27me3), separated by state change characterisation.

**Figure S6.**
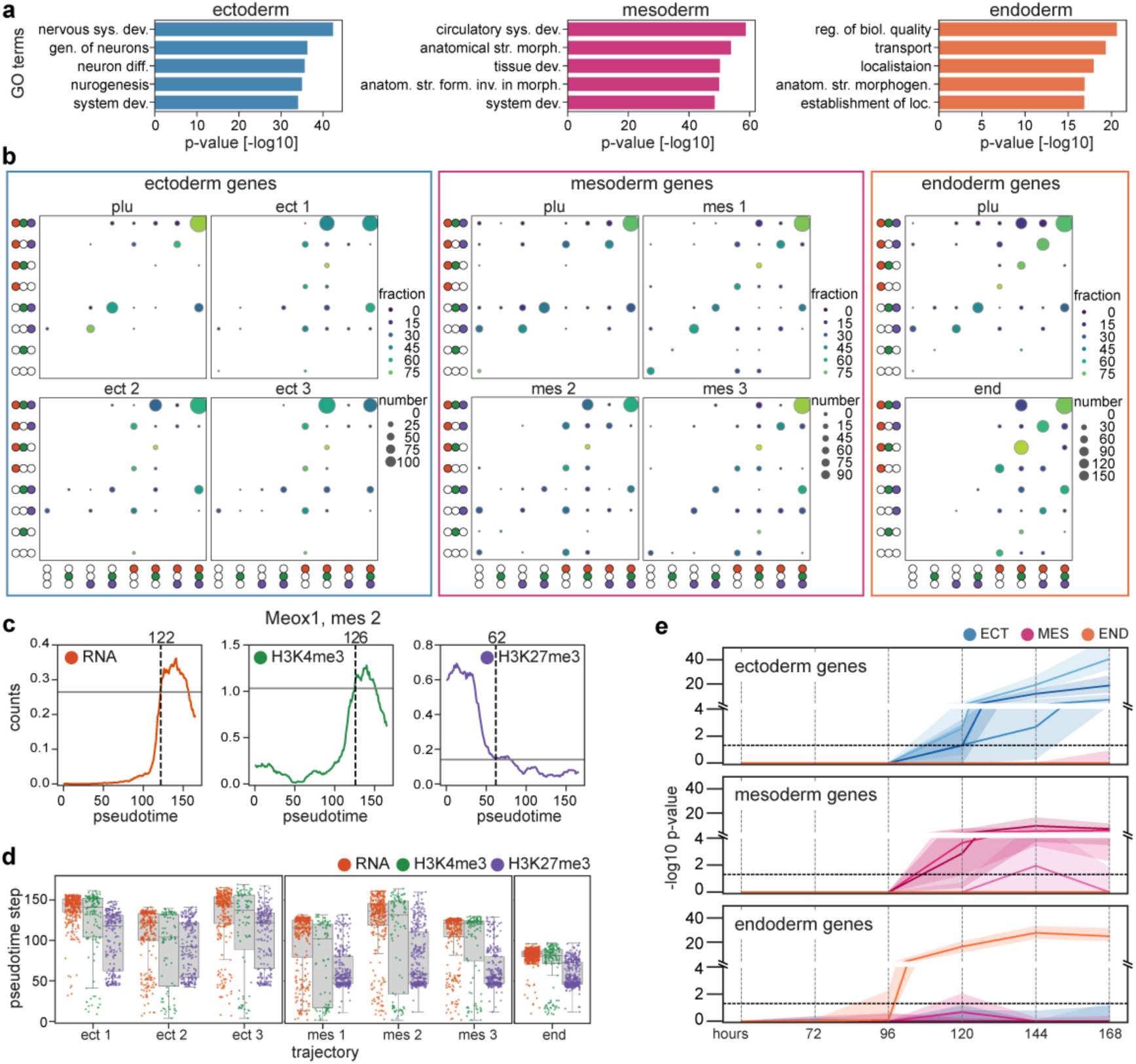
Coordination between transcriptional activation and H3K27me3 differs between TFs and their targets. a) GO term analysis of differentially expressed genes for the three germ layers (left: ectoderm, middle: mesoderm, right: endoderm). b) Summary of state changes of differentially expressed genes for ectoderm (left), mesoderm (middle) and endoderm (right) gene sets in the ESC to pluripotent as well as their respective trajectories are shown. Dot size indicates number of genes per state change, colour indicates fraction of genes with the same start state changing to indicated state. c) Indication of 2x standard-deviation threshold in the three modalities for example gene Crabp2 in the ectoderm 3 trajectory. Grey horizontal lines indicate standard deviation threshold, dotted vertical line indicates the first identified step of significant change. d) Boxplot summarising at which step along the trajectory the differentially expressed gene sets change for RNA (orange), H3K4me3 (green), and H3K27me3 (purple) in all trajectories. e) Line plots indicating the -log10 p values of differentially expressed gene sets relative to mESCs over sampling time (top: ectoderm-specific genes, middle: mesoderm-specific genes, bottom: endoderm-specific genes). The 7 trajectories are indicated as lines: ectoderm: light to dark blue, mesoderm: light to dark pink, endoderm: orange. Dashed line indicates significance threshold.

